# Safety profiling of genetically engineered Pim-1 kinase overexpression for oncogenicity risk in human c-kit+ cardiac interstitial cells

**DOI:** 10.1101/521732

**Authors:** Kathleen Broughton, Kelli Korski, Oscar Echeagaray, Robert Adamson, Walter Dembitsky, Zhibing Lu, Erik Schaefer, Mark A. Sussman

## Abstract

Advanced approaches to stem cell-based therapies is necessary for myocardial regenerative therapy because treatments have yielded modest results in the clinic. Our group previously demonstrated genetic modification of cardiac stem cells with Pim-1 kinase overexpression rejuvenated aged cells and potentiated myocardial repair. Despite these encouraging findings, concerns were raised regarding oncogenic risk associated with Pim-1 kinase overexpression. Testing of these c-kit+ cardiac interstitial cells (cCICs), derived from heart failure patient samples, overexpressing Pim-1 (cCICs-Pim-1) for indices of oncogenic risk was assessed by soft agar colony formation, micronucleation, gamma-Histone 2AX foci, and transcriptome profiling. Collectively, findings demonstrate comparable phenotypic and biological properties of cCICsPim-1 compared to baseline control cCICs with no evidence for oncogenic phenotype. Using a highly-selective and continuous sensor for quantitative assessment of PIM1 kinase activity, a 7-fold increase in cCICs-Pim-1 versus cCICs resulted. Kinase activity was elevated in IKKs, AKT/SGK, CDK1-3, p38, and ERK1/2 in addition to Pim-1, correlating Pim-1 overexpression to contribute to Pim-1-mediated effects. Enhancement of cellular survival, proliferation, and other beneficial properties to augment stem cell-mediated repair without oncogenic risk is a feasible, logical, and safe approach to improve efficacy and overcome current limitations inherent to cellular adoptive transfer therapeutic interventions.

## Introduction

Stem cell-based therapeutic approaches are poised to revolutionize treatment of numerous diseases previously considered untreatable ^1-10^, but further improvements and enhanced products are needed to make this therapy truly safe and efficacious. Longstanding challenges of poor survival for adoptively transferred cells and low engraftment rates have stymied progress toward enhancing efficacy, especially in the area of myocardial regenerative medicine.^11-17^ Our group has pioneered use of genetic engineering to enhance regenerative potential of human c-kit+ Cardiac Interstitial Cells (cCICs), historically referred to as cardiac stem/progenitor cells (CPCs), intended for autologous therapeutic utilization. cCICs support myocardial repair and improve cardiac performance through reduction of scar size and increased cardiac output.^18^ Pim-1, an endogenous constitutively activated enzyme found within cCICs, is produced in response to stress or pathologic injury in the myocardium.^19^ Pim-1 up-regulation enhances cell survival through mediating transcription, cell growth, proliferation, survival, and expansion of cCICs^20,21^. Our previous efforts with over a decade of research on Pim-1 biology demonstrates remarkable regenerative effects mediated by adoptively transferred human Pim-1 enhanced CPCs in rodent and swine models with significantly improved cardiac performance relative to unmodified CPCs, bone marrow stem cells and mesenchymal stem cells.^22-24^ Human Pim-1 enhancement of CPCs also exhibits reversal of aging affects, such as down regulation of senescence markers, increased telomere length and mitochondrial activity and the augmented proliferation.^25^ The technology of using autologous Pim-1 enhanced cCICs (cCICs-Pim-1), patented as CardioEnhancers™, represents convergent evolution of extensive high-level mechanistic biological research that creates a novel and efficacious treatment for heart failure.^26^ However, association of Pim-1 as a marker of oncogenic potential^27-29^ has raised some concerns regarding the potential for transgene-mediated overexpression of Pim-1 to promote increased oncogenic risk using cCICs-Pim-1. ^30^ Furthermore, concerns continue to persist regarding insertional mutagenesis through use of lentiviral vectors for genetic modification that could promote oncogenic transformation.^31-35^ Establishing the safety profile of cCICs-Pim-1 regarding oncogenic risk is a recurring recommendation to advance implementation, forming the rationale for studies performed in this report.

Pim-1 protein is an evolutionarily highly conserved serine / threonine kinase unique by virtue of constitutive activation, meaning that rather than having to be ‘activated’ as with most kinases, it is active in nascent translated form.^36^ Thus, Pim-1 activity is regulated by concerted control of gene transcription, mRNA translation, and protein degradation. The target phosphorylation consensus sequence for Pim-1 is found in proteins mediating transcription, cell growth, proliferation, and survival.^37,38^ Transgenic mouse experiments^39-41^ and cell culture experiments^42-45^ demonstrate PIM-1 exerts potent synergistic activity with c-MYC, p21Cip1/WAF1, STAT3, JNK, and survivin, as well as with other proteins particularly when the protein function involves proliferation and cell survival. However, it is very important to recognize that PIM-1 is considered a weak oncogene at best because it is not a transforming factor by itself and this assessment has not significantly changed over the years. Murine myeloid cells in culture are dependent upon IL-3 and rapidly die if the IL-3 was withdrawn, suggesting a major function of Pim-1 is to act as a survival factor.

Further support of this notion comes from the observation that forced expression of PIM-1 improves factor-dependent cells survival in the absence of IL-3^46^. Pim-1’s role in cell survival led to the discovery that it down regulates activity of the ASK1 proapoptotic pathway.^47^ In general, Pim-1 upregulation enhances cell survival whereas loss of Pim-1 increases apoptotic cell death. Protective effects of Pim-1 depend upon kinase activity as borne out by experiments using a dominant-negative kinase dead mutant construct.^46,48^ In fact, Pim-1 transgenic overexpression in lymphoid tissue showed “Eµ-pim-1 mice lack sufficient sensitivity to justify their routine use in short-term carcinogenicity testing in general.” Despite increased predisposition to tumor induction in the lymphoid compartment, Pim-1 overexpression did not increase susceptibility to oncogenic induction by genotoxic carcinogens.^49^ Protein kinases linked to cancer development may share phosphorylation consensus sequences with Pim-1, but their modes of regulation are dissimilar, their expression patterns differ, and separate signal transduction pathways activate them. The collective complexities of Pim-1 biology, lentiviral genetic engineering, and uncertainties regarding oncogenic risk justified a thorough and comprehensive study to profile the effect of Pim-1 overexpression in human cCICs.

## Materials and Methods

### Cell Culture

K562 cells, a human bone marrow lymphoblast cell line derived from a female patient with chronic myelogenous leukemia (CML), were obtained from ATCC^®^ (CCL-243^(tm)^). Human cCICs cell lines for individual isolates from three patients undergoing surgical implantation of a left ventricular assist device (LVAD) were used for characterizations. cCICs were isolated and expanded as previously described^50^ and each was subjected to lentiviral modification to express Pim-1 GFP in a bicistronic construct or GFP alone as a control cell line. Culture media formulations used for cell culture and assays are presented in Supplemental Table 1. Cell pellets (∼1 x 10**^6^** cells washed with 1x PBS) were prepared and snap frozen in liquid nitrogen (or dry ice) and stored at −80 **°**C for up to 3 months before use as described below.

### Lentiviral Infection

Lentiviral preparations were created as previously described^23,24^ with modifications as detailed in Supplemental Methods section. Early passage (4-5) cCICs were plated over night at density of 50,000 cells/well on a 6-well plate. The following day cells were washed with 1xPBS prior to adding 1mL of cCICs medium (Supplemental Table 2). Lentivirus infection was performed at a multiplicity of infection ratio of 3 particles per cell, with virus added per well (PIM1 L1701 titer 2.6 x 10^8^ TU/mL; GFP L1702 titer 1.8 x 10^8^ TU/mL) in 1mL of warm cell culture medium. The plate was intermediately shaken throughout the day. At the end of the day, another mL of warm cell culture medium was added. The next morning the virus removed, cells were washed with PBS and replenished with cell culture medium. 2 days after infection cells were divided into two T25 flasks. 2 days after the division, one of the flasks was continued for expansion and the other one was used for FACS and protein quantification.

### Immunoblotting

Cells were plated 24-hours prior to lysis (50,000 cells/well 6-well plate). On the day of collection, cells were washed with cold 1xPBS, placed on ice and lysed with 50µL of sample buffer with protease and phosphatase inhibitors (added immediately before use), followed by boiling for 5 minutes and sonication. Lysates were cleared by centrifugation before loading on to 4%-12% NuPage Novex Bis Tris gel (Invitrogen), transferred on to a polyvinylidene fluoride (PVDF) membrane and blocked for an hour at room temperature in 5% skim milk, 1x tris-buffered saline tween-20 (TBST). Primary antibodies were incubated over night at 4°C. PIM-1 (Thermo, ZP003) 1:1,000; GFP (Rockland, 600-101-215) 1:500; GAPDH (Sicgen, AB0067-200) 1:1,000 in 5% skim milk. Membranes were washed with TBST 3×5 minutes. Secondary antibodies (Licor, 1:2,000) were incubated for an hour at RT, followed by 3×5 minutes TBST wash. Membranes were imaged on Licor Odyssey CLx.

### Immunocytochemistry

Next day after plating 40,000 cells/well in a glass bottom chamber slide were fixed in 4% formaldehyde (Thermo, 28908) for 5 minutes, rinsed with 1xPBS 3×5 minutes. Cells were permeabilized with 0.1M Glycine for 5 minutes followed by 0.1% Triton-x100 treatment for 10 minutes at RT. Followed by 3×5 minute wash in 1xPBS. Cells were blocked using TNB blocking buffer (0.1M Tris-HCl, pH 7.5; 0.15M NaCl; 0.5% TSA blocking reagent (Perkin Elmer, FP1012) for 1 hour at RT. Primary antibody (yH2AX (abcam) 1:200) was incubated over night at 4°C in TNB. After incubation cells were washed 3×5 minutes 1xPBS before secondary antibody (1:200) and phalloidin 488 (1:200) incubation (1hr, RT). Cells were washed with 3×5 minutes of 1xPBS, where DAPI was included in the middle wash at 1:1,000. Slides were coverslipped and imaged on SP8 Leica confocal microscope.

### Soft Agar Assay

Bottom layer of 0.5% noble agar (Sigma, A5431) in either cCIC or HEK293 cell culture medium (Supplemental Table 1) with a 1x final concentration was plated on 6-well plates (1mL/well) in sterile conditions. The agar was allowed to solidify under laminar flow for at least 30 minutes. Meanwhile, the cells were lifted, counted and a cell suspension of 6,667 cells/mL was prepared in 2x cell culture medium. Cells were mixed 1:1 with 0.6% noble agar solution at 42°C to avoid immature solidification of agar. 1.5mL of cell solution (5,000 cells/well) in agar was plated on top of 0.5% agar base. The cell and agar mixture solidified after 30 minutes under a laminar flow hood prior to placing into a cell culture incubator. To maintain sufficient humidity and nutrition levels in the wells, 100µL of cell culture medium was added twice a week. After 14 days 200µL of nitroblue tetrazolium (Invitrogen, N6495) was added to the wells and incubated overnight in a cell culture incubator. The following day the wells were scanned using Licor Odyssey CLx and the colonies were counted using ImageJ software. A more detailed protocol has previously been published^1^.

### In Vitro Micronucleus Assay

On the first day 50,000 cCICs were plated per well on a 6-well plate. The following day, the cells were washed with 1mL of 1xPBS prior to adding 4µg/mL of cytokinesis block cytochalasin B (Sigma, C2743-200UL) for 24 hours. The working concentration of cytochalasin B was experimentally decided by finding the concentration with least observable cytotoxic effects but the highest number of double-nucleated cells. After the 24-hour treatment, the supernatant was collected, cells lifted and pelleted. The pellet was re-suspended in 400µL of 1xPBS, 0.5% BSA and placed on ice. 200µL of cell suspension was loaded onto cytofunnels (Sigma, A78710020), the funnels were spun on Thermo Shandon Cytospin 4 at a speed of 400 rpm for 3 minutes on low acceleration. The final cell number 25,000 cells/slide. Immediately after cytospinning, the cells were fixed with 4% formaldehyde (Thermo, 28908) for 5 minutes, followed by 3 x 5-minute washes of 1xPBS. The cells were stained with DAPI (1:1,000) and FITC conjugated phalloidin (1:200) (Invitrogen, A12379) for 30 minutes in TNB blocking buffer (0.1M Tris-HCl, pH 7.5; 0.15M NaCl; 0.5% TSA blocking reagent (Perkin Elmer, FP1012)). Cells were washed with 3 x 5 minutes of 1xPBS, cover slipped with Vectashield (Vector, H-1000) prior to analysis on Leica SP8 confocal microscope. Z-stack images were taken with 40x water objective of each sample set. Only the double-nucleated cells, with clear cell borders were included in the analysis for micronuclei. The analysis included 3 biological replicates, for each cell line at least 667 cells were counted, with a total number of 2,000 cells per experimental condition. Data is represented as percent micronuclei per biological replicated with standard deviation. Student’s t test was used to calculate significance between the two groups. The protocol is adjusted from the OECD/OCDE guidelines for the testing of chemicals.^2^

### Bulk RNA-Seq

Total RNA was extracted using Quick-RNA Miniprep Kit (Zymo, R1054) as per manufacturer’s instructions. RNA concentration and purity were assessed by Nanodrop and RNA integrity was assessed using the Agilent TapeStation. cDNA Libraries were prepared for mRNA-enriched sequencing using TruSeq Stranded mRNA kit (Illumina). This was followed by normalization to 6 nM and pooling of libraries, followed by single end 75 bp sequencing on the Illumina HighSeq 4000. Reads were aligned to the GRCh38.24 genome using HiSat2, transcripts were assembled and expression frequencies were assessed from the aligned data by StringTie, and Ballgown 2.12.0 was used to identify DEGs and generate the gene expression values and plotting.^51^ FASTA Sequence was used for downstream bioinformatic tracking of PIM-1 splice forms (provided in Supplemental Methods).

### Bioinformatic statistics

Differential expression analysis was performed using R statistical package Ballgown 2.12 using a previously described standard linear model-based comparison^1^. The threshold of selection of differentially expressed genes was the following: gene abundance of FPKM>1, p value < 0.05 and a log2 fold change of ±1. Significant differences in the number of detected genes and ratios between expressed and differentially expressed genes were analyzed using the non-parametric t-test Mann-Whitney, with statistical significance accepted of p < 0.05.

### Data availability

RNA-Seq data generated in this study has been uploaded to the Gene Expression Omnibus (GEO) database (GSE124590).

### Kinase Activity Measurements

Kinase activity was measured continuously using the PhosphoSens® technology (AssayQuant Technologies Inc., Marlborough, MA) according to the manufacturers’ recommendations. This one-step homogeneous assay format uses chelation-enhanced fluorescence via optimized substrate sensors containing the unnatural fluorogenic amino-acid Sox.^52-58^

### Recombinant Kinases

The following kinases were obtained from Carna Bioscience and used at 1 nM in this study: PIM1 (amino acids 1-313, cat. & lot #: 02-054, 10CBS-0421F), PIM2 (amino acids 1-311, cat. & lot #: 02-155, 10CBS-0283E), PIM3 (amino acids 1-326, cat. & lot #: 02-156, 12CBS-0631B), p70S6 (amino acids 1-421, cat. & lot #: 01-156, 14CBS-0617B), RSK1 (amino acids 1-735, cat. & lot #: 01-149, 08CBS-0354K), CHEK2 (amino acids 1-543, cat. & lot #: 02-162, 10CBS-0386B), AKT3 (amino acids 108-479, cat. & lot #: 01-103, 09CBS-1278 E) and MAPKAPK2 (amino acids 1-400, cat. & lot #: 02-142).

### Crude Lysate Preparation

Cells pellets stored at −80 ° C were thawed on ice prior to resuspending in Cell Extraction Buffer (CEB) containing 50 mM TRIS, pH 7.5, 150 mM NaCl, 2 mM EGTA, 30 mM NaF, 10 mM Na_4_P_2_O_7_, 100 μM Na_3_VO_4_, 1% Triton X-100, 50 mM β-glycerophosphate, 1 mM DTT, Sigma Protease Inhibitor cocktail (#P8340) 10%; Sigma Phosphatase Inhibitor cocktail (#P285) 10%. Specifically, cell pellets from His067/P-12/PIM-1 or His067/P-12/GFP control conditions (1×10^6^ cells) were solubilized in 0.6 ml CEB and sonicated on ice for 2 seconds X10 with 5 seconds cooling in between sonication. Clear cell lysate was obtained after centrifuging @6000g for 10 minutes @ 4 C. Control samples of K562 cells, which are known to exhibit Pim-1 kinase activity, were generated as described above to serve as positive controls for the assay. The Bradford method was used to determine the total protein concentration of clear cell lysate, which was ∼ 1 mg/mL for all three samples. Once generated, crude cell lysate samples were kept on ice and immediately assayed for kinase activity using a panel of CSx-Substrates as described below.

### Final Reaction Conditions

Protein kinase activity was determined in 54 mM HEPES, pH 7.5, 1 mM ATP, 1.2 mM DTT, 0.012% Brij-35, 0.52 mM EGTA, 10 mM MgCl_2_, 1% glycerol, 0.2 mg/ml BSA, and 10 μM Sox-based sensor. A panel of Sox-based sensors (AQTx) was used to measure the kinase activity of tyrosine kinases (AQT0001) or serine/threonine kinases including PIM1(AQT572181C09), CK1 (AQTST24), IKK-family (AQT0215), AKT and SGK family (AQT0233), CDK1-3 and 5 (AQT0255), CDK4 and 6 (AQT0258), JNK/MAPK (AQT0365), p38/MAPK (AQT0374), and ERK1/2 MAPK (AQT0376). All components except enzyme were equilibrated to 30°C prior to setting up reactions in 96-well half-area, white, flat and round-bottom NBS microplates (Corning #3824). For each reaction (50 μL final volume), 5 μL Sox-based substrate (10x) was first mixed with35 μL Reaction Mix containing ATP & DTT (1.43x), followed by a 5-minute preincubation (all components except buffer control or cell lysate) at 30°C. This was followed by addition of 10 μL CEB (1x) or crude cell lysate (5x in CEB) or 1-2.5 μg of crude cell lysate per reaction. Actual final reaction concentrations containing CEB were 10 mM TRIS, pH 7.5, 30 mM NaCl, 6 mM NaF, 2 mM Na_4_P_2_O_7_, 20 μM Na_3_VO_4_, 0.2% Triton X-100, 10 mM β-glycerophosphate, Sigma Protease Inhibitor cocktail (#P8340) 2%, and Sigma Phosphatase Inhibitor cocktail (#P285) 2%). Plates were sealed using optically-clear adhesive film (TopSealA-Plus plate seal, PerkinElmer, applied with a roller) to eliminate evaporation and resulting drift and then fluorescence intensity measurements were read kinetically every 2 minutes from the top for up to 120 minutes at 30°C, with excitation and emission wavelengths of 360 nm and 485 nm, respectively, using a Synergy Neo2 multi-mode plate reader (Biotek Instruments, Winooski, VT).

### Kinase Activity Analysis

Fluorescence, determined with identical reactions but lacking purified enzyme or crude cell lysate was subtracted from the total fluorescence signal for each time point, with both determined in duplicate, to obtain corrected relative fluorescence units (RFU). Corrected RFU values then were plotted versus time and the reaction velocity for the first ∼40 minutes (initial reaction rates) were determined from the slope with units of RFU/min or RFU/min/μg of crude cell lysate.

### Statistical Analysis

Data are presented as mean ± standard deviation (SD). For comparisons of two groups, Student’s t-test was used to determine significance. For comparisons of multiple groups, one-way ANOVA was performed followed by Tukey’s post hoc test. Statistics were performed using GraphPad Prism (LaJolla, CA) software.

## Results

### Lentiviral engineering of human cCICs for stable overexpression of Pim-1 kinase

Cell lines derived from three individual heart failure patient samples (see Table 1) were identified as Line #1 (H15-096), Line #2 (H13-067), or Line #3 (H16-107). The cCIC lines expanded from each patient sample were genetically engineered using lentiviral vectors expressing either GFP alone or GFP and Pim-1 in a bicistronic construct. Efficiency of genetic modification of patient cell lines was assessed by flow cytometric analyses (Figure 1A). GFP expression alone in stably modified cCICs was 78.5%, 88.5%, and 80.2% for Lines 1, 2, and 3, respectively. Similar efficiency of GFP expression in cCICs-Pim-1 for expression of GFP / Pim-1 was observed at 81.7%, 84.7%, and 80.9% for Lines 1, 2, and 3, respectively. Protein expression of GFP as well as Pim-1 in cell lysates from each line was confirmed by immunoblot analysis (Figure 1B), with an average significant increase of Pim-1 kinase expression in engineered lines of 25-fold increase in GFP / Pim-1 cCICs relative to corresponding cells expressing GFP alone (Figure 1C). These results demonstrate the efficient and consistent genetic modification of human cCICs to overexpress Pim-1 at levels significantly higher than normally found in these cells.

**Table 1.** Patient Data

| Nr | Line | Gender | Age | ischemic | Diabetes | Smoker |
| --- | --- | --- | --- | --- | --- | --- |
| 1. | H13-067 | male | 84 | MI + CABG | + | quit 1991 |
| 2. | H15-096 | male | 68 | ischemic + CABG | N/A | N/A |
| 3. | H16-107 | male | 71 | ischemic | N/A | past smoker |

**Figure 1.**
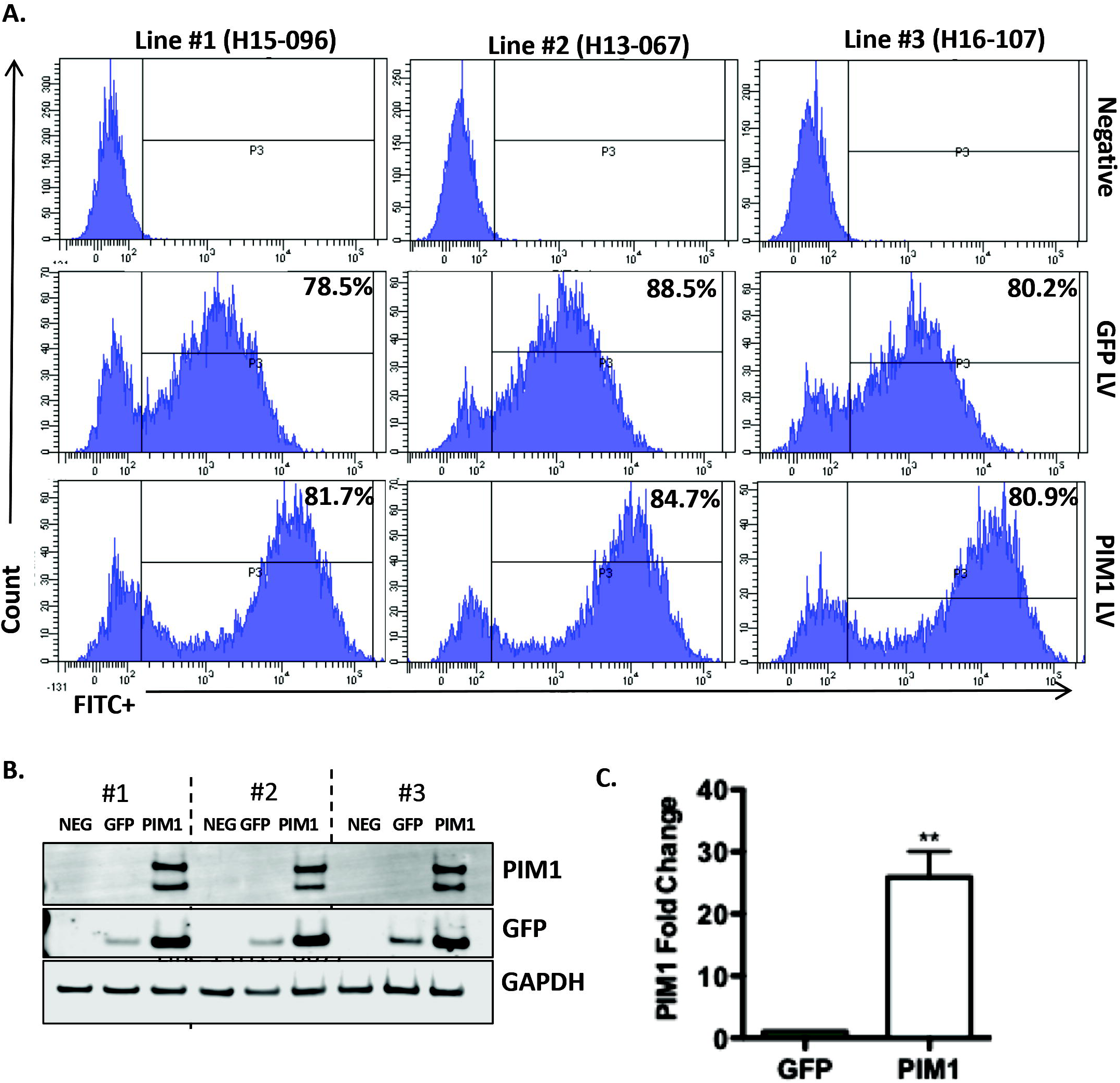
PIM1 kinase over expression via lentiviral vector in human cCICs. (A) FACS derived histograms representative of three human cCIC lines infected with either GFP lentivirus or Pim1 lentivirus (MOI 3). Histograms are gated relative to passage matched non-infected control. (B) Immunoblot of three lines #1(H15-096), #2(H13-067), #3(H16-107). (C) Quantification of Pim1 levels in Pim1 O/E samples relative to GFP O/E samples. Normalized to GAPDH. Paired student’s t test. N=3 P^**^<0.01

### Soft agar colony growth is not induced in Pim-1 overexpressing cCICs

Anchorage-independent growth in soft agar is a well described assay for oncogenic transformation of cells.^59,60^ Ability of cCICs to proliferate in soft agar was assessed for lines expressing GFP alone as well as GFP / Pim-1 (Figure 2). The positive control cell line HEK293 demonstrated growth competency in the soft agar assay, forming colonies clearly visible within 2 weeks after plating (Supplemental Figure 1). In contrast, cCICs lines were unable to form colonies in soft agar and were all scored as negative for growth at the two-week time point. All three cCIC lines showed comparable negative results, with representative images shown for Lines 1 and 3 (Figure 2). These results demonstrate that Pim-1 overexpression in cCICs fails to promote growth of the cells in soft agar consistent with normal, non-transformed cell phenotypic properties.

**Figure 2.**
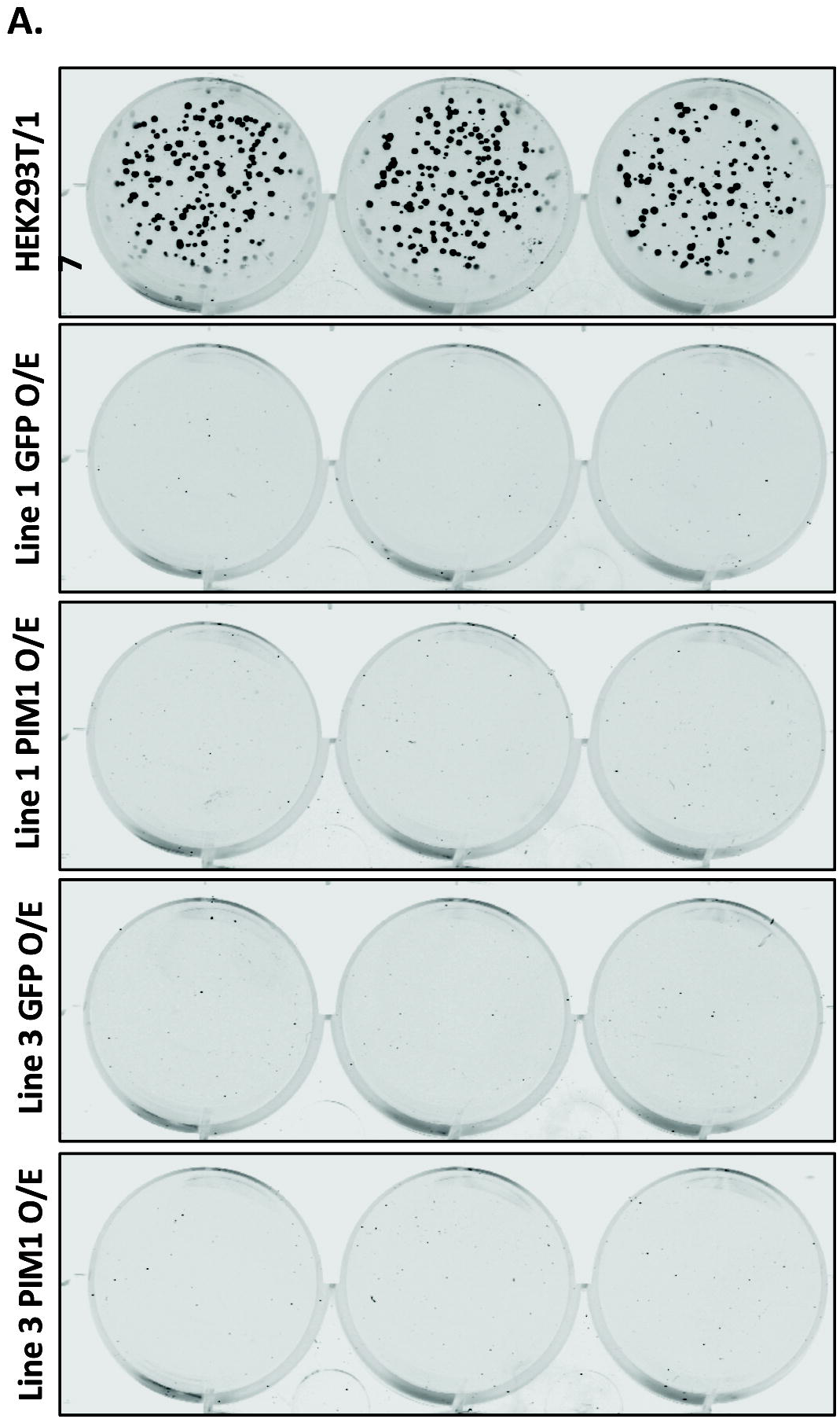
Soft Agar Assay. (A) Colony formation after 2 weeks (5,000 cells/well). Positive control HEK293T/17 (passage 7) all samples are in triplicates. N=2 From top to bottom: Line 1(H15-096) GFP O/E and PIM1 O/E, Line 3 (H16-107) GFP O/E, PIM1 O/E.

### Micronucleation is not increased in Pim-1 overexpressing cCICs

Standardization of protocols and readouts for *in vitro* micronucleation transformation (IVMNT) over decades of use has resulted in a reliable, robust, and straightforward method for presence of micronuclei as a marker of chromosome damage. Cell lines of cCICs derived from expansion were prepared for IVMNT using established protocols^61-66^ with cytochalasin B treatment to promote binucleation (Supplemental Figure 2). Control cells include lentivirally-modified eGFP expressing cells without Pim-1 as well as HEK293T/17 cells as positive controls for clastogenic and aneugic events (Supplemental Figure 3). Presence of micronuclei indicative of genotoxic damage was quantitated using microscopic assessments (Figure 3). Micronuclei formation was comparable between all three cCIC lines engineered with GFP alone versus cell expressing GFP / Pim-1 without statistical significance (p=0.5). These results demonstrate that overexpression of Pim-1 in cCICs does not promote micronucleation indicative of chromosome instability and potential oncogenic transformation.

**Figure 3.**
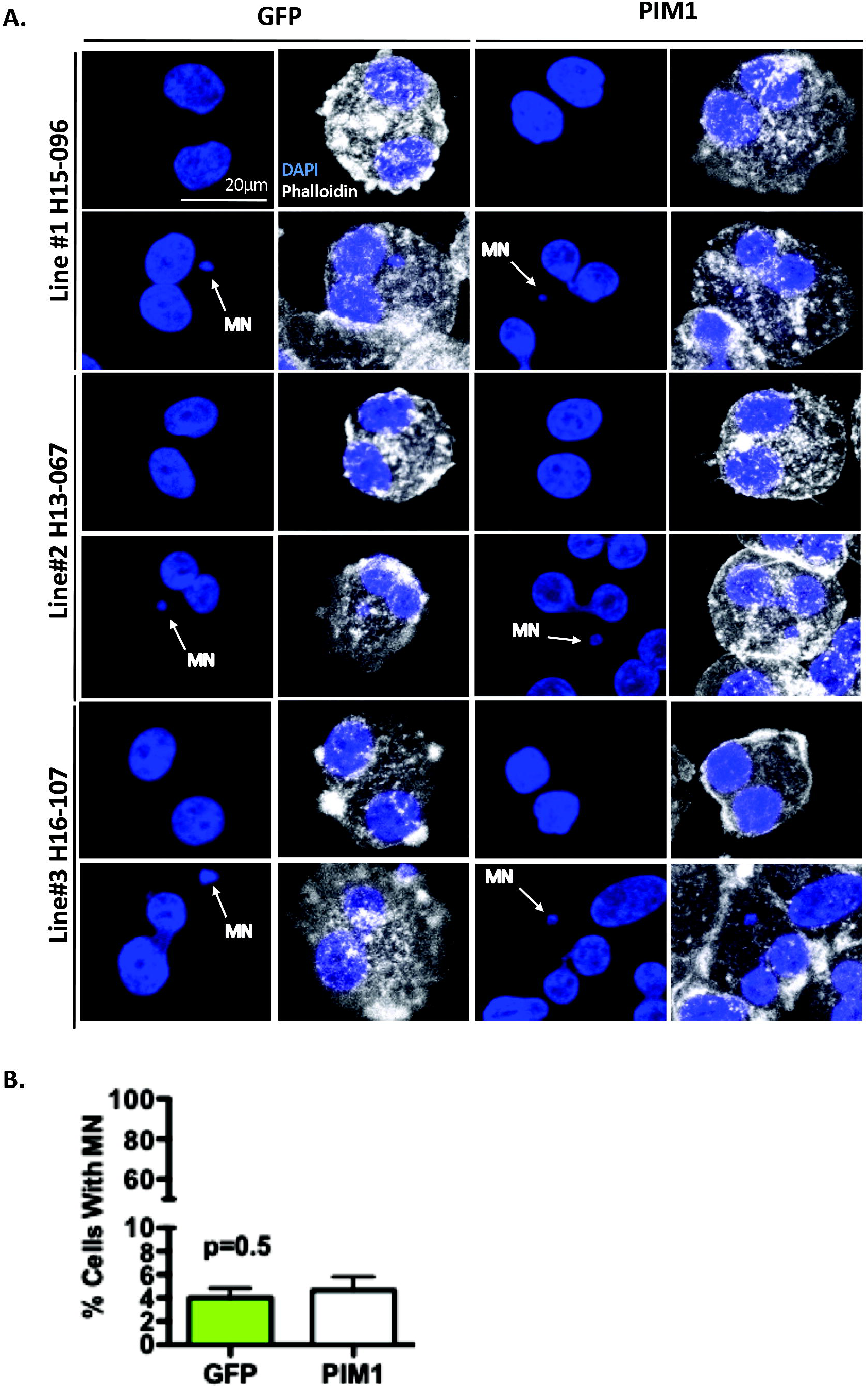
PIM1 Over Expression Does Not Significantly Increase Micronucleation. (A) Representative images of cells with normal nuclei and cells with micronuclei for each cell line, with and without PIM1 O/E. Cells were treated with 4µg/mL cytochalasin B to increase the number of binucleated cells. Micronuclei (MN) are pointed out by white arrows. DAPI (blue), Phalloidin (white). 40x water objective, scale bar 20µm. (B) Quantification of percent MN in binucleated cells in lines #1 (GFP 682 cells, 22 MN; PIM1 668, 30 MN), #2 (GFP 717 cells, 35 MN; PIM1 691 cells, 36 MN), #3 (GFP 669 cells, 22 MN; PIM1 674 cells, 22 MN). Total # cells analyzed for GFP 2,068 (79 MN) and PIM1 2,033 (88 MN). N=3 Paired student’s T test.

### Foci of γ-H2AX immunolabeling are not increased in Pim-1 overexpressing

**cCICs.** DNA double-strand breaks and blocked replication forks are marked by phosphorylation of histone 2AX (H2AX) that accumulates at the site of damage and can be visualized as foci by immunocytochemistry. The three cCIC lines were prepared for immunofluorescence confocal microscopy and immunolabeled for γH2A.X (phospho-S139) with DAPI as a nuclear counterstain (Figure 4A). Control cells were lentivirally-modified Lines expressing GFP alone without Pim-1. Presence of punctate nuclear-localized labeling characteristic of yH2AX (phospho-S139) immunolabeling indicative of genotoxic damage was quantitated using microscopic assessments of 50 cells per experimental group. Number of γH2A.X foci per nucleus were comparable between all Lines tested and no statistically significant difference was observed between Lines expressing GFP alone versus those expressing GFP / Pim-1 (Figure 4B; p>0.5). These results demonstrate that overexpression of Pim-1 in cCICs does not promote formation of γH2A.X (phospho-S139) foci indicative of DNA damage.

**Figure 4.**
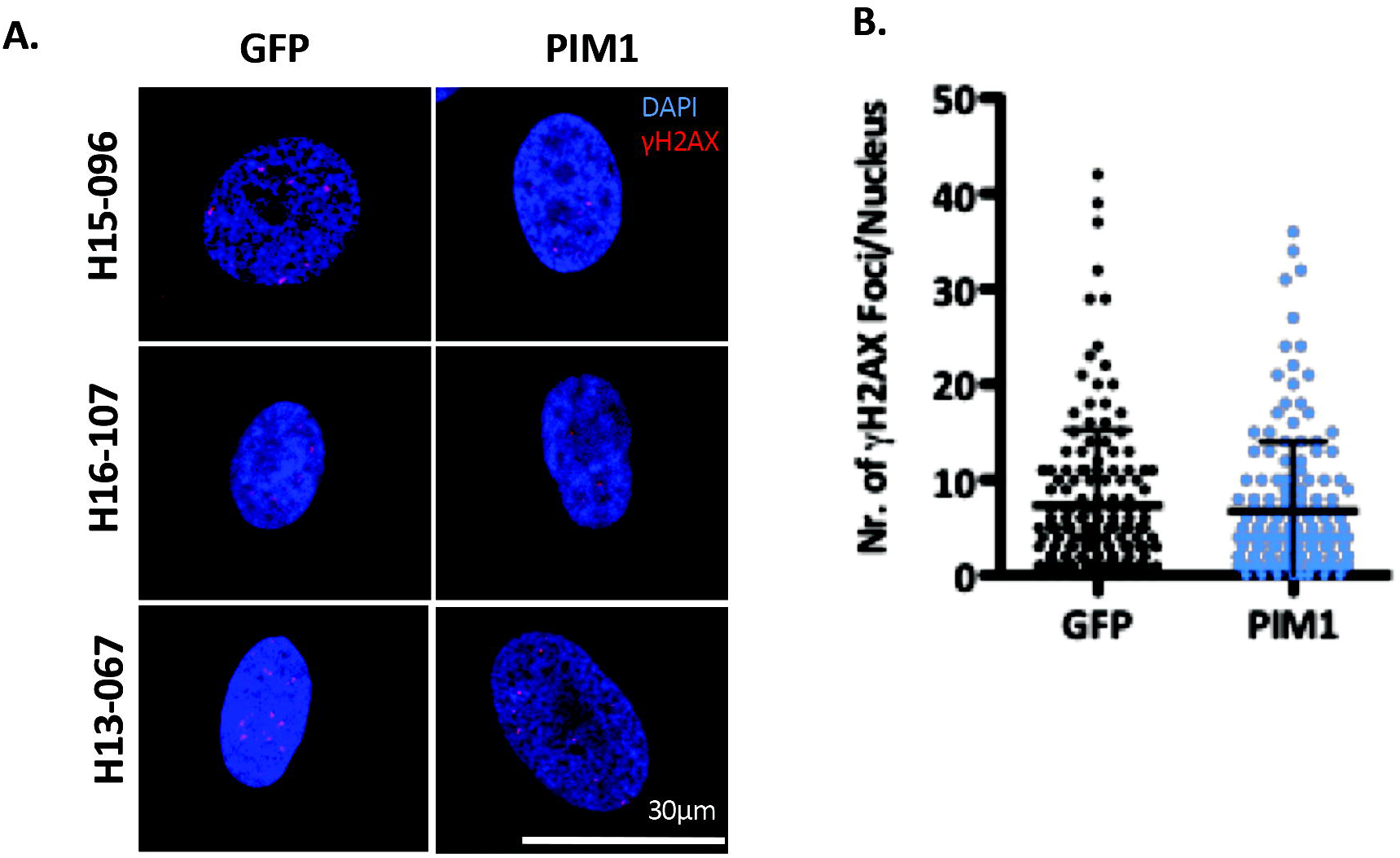
γH2AX in human cCICs. (A) ICC Images of cCICs stained with γH2AX (red), DAPI (blue). 40x water objective. Scale bar 30µm. (B) Quantification of γH2AX foci per nucleus in GFP O/E and PIM1 O/E samples. (50 cells per cell type, per condition). Student’s t test. N=3 p=0.5.

### Transcriptome profiles are not significantly altered by Pim-1 overexpression in cCICs

Phenotypic characteristics of cCICs show no evidence for increased oncogenic potential from Pim-1 overexpression (Figures 2-4), prompting further studies to assess other aspects of altered biological properties related to oncogenic markers. Transcriptional profiling by bulk RNA-Seq analysis was performed, choosing a mixture of Lines 1, 2, and 3 as a representative sample since all three lines exhibited comparability for expression of Pim-1 (Figure 1) and similarly negligible oncogenic hallmarks (Figures 2-4). Correlation analysis of gene expression profiles demonstrated that transcriptional profiles were not significantly altered between cCICs expressing GFP alone versus cell expressing GFP / Pim-1 (Fig. 5A). Furthermore, transcriptional impact of Pim-1 overexpression is minimal compared to inherent patient transcriptional diversity (Fig. 5B). A total of 25 differentially expressed genes (DEGs) were detected with only 14 upregulated genes in Pim-1 overexpressing cCICs (Fig. 5C). Individual and collective quantification of normalized transcripts and subsequent alignment to spliced isoforms was performed to confirm overexpression of Pim-1 variant 1 despite minimal differences in the global transcriptome (Supplemental Figures 4 and 5, confirmatory FASTA sequences of overexpressed splice form of PIM-1). Additionally, no significant differences were observed in the overall detected genes, genes displaying high abundance (FPKM>1), or the relationship between detected genes and DEGs (Fig. 5D and Supplemental Figure 6). Collectively, these results confirm lack of significant transcriptional reprogramming pursuant to Pim-1 overexpression in cCICs.

**Figure 5.**
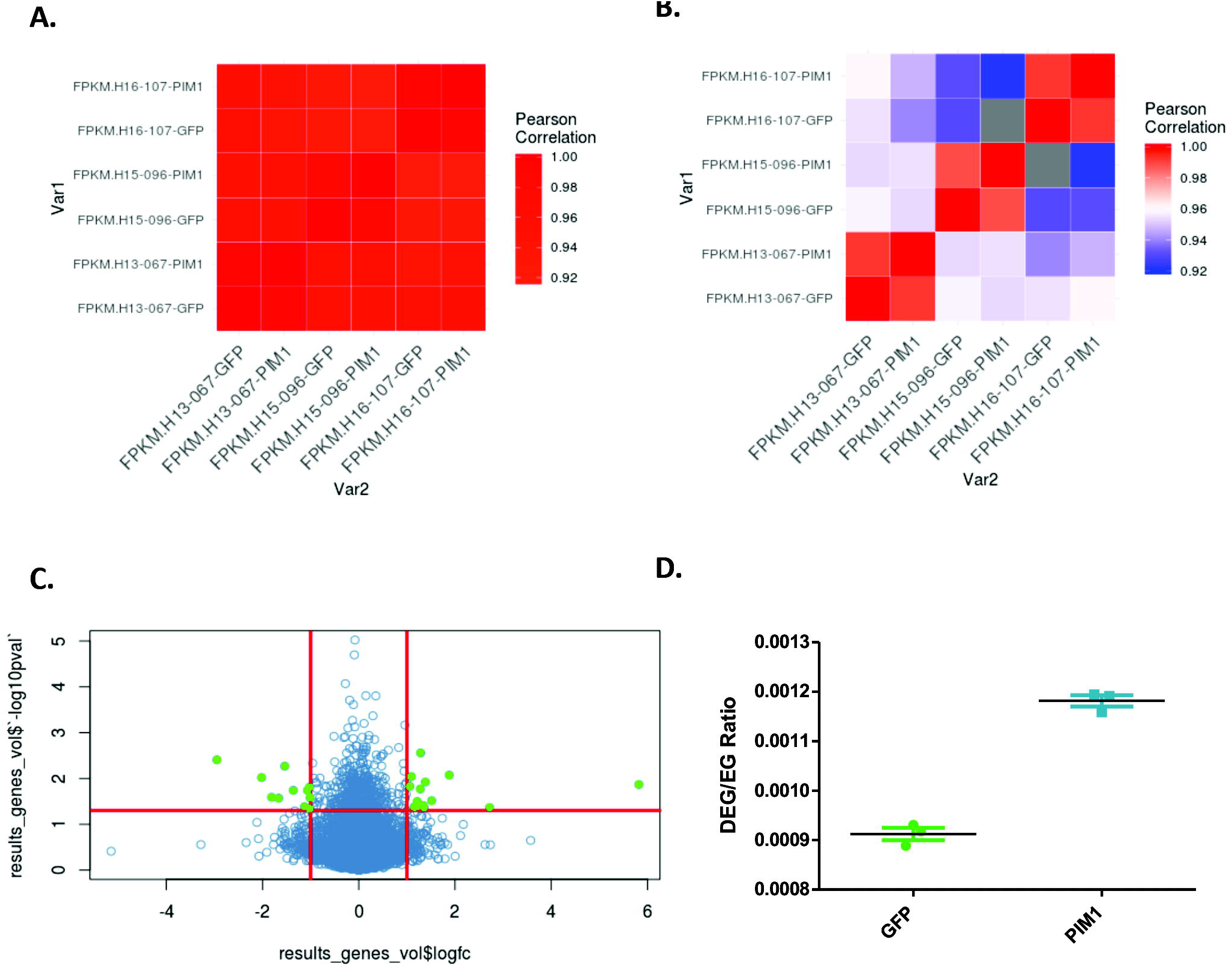
PIM-1 enhancement minimally impacts the transcriptome of cCICs-Pim-1 revealed by RNA-SEQ. Pearson’s correlation analysis between PIM-1 enhanced and transfection control of total expressed genes displaying (A) high transcriptional similarity (correlation coefficient r^2^>0.92) and (B) a transcriptional impact of PIM enhancement below the inherent patient diversity. (C) Volcano plot displaying 25 significantly differentially expressed genes of PIM enhanced cell lines compare to control (p<0.05, log2(Fold Change) >±1). (D) PIM enhancement does not significantly impact the ratio of differentially expressed genes to total expressed genes (Mann-Whitney t-test, p=0.1).

### Kinase profiling using real-time sensors reveals elevation of PIM-1 activity and multiple Pim-1 target pathways in cCICs

Given the critical role of PIM-1 activity in conferring the beneficial properties of cCICs, we felt it was very important to measure PIM-1 activity directly. The Imperiali laboratory has pioneered the use chelation-enhanced fluorescence via optimized substrate sensors containing the unnatural fluorogenic amino-acid Sox.^52,55-57^ This one-step homogeneous assay format provides a direct, continuous (kinetic) and highly-quantitative measure of kinase activity that can be performed with purified enzyme or crude cell or tissue lysates.^53,54,58,67^ Using a highly-selective sensor for quantitative assessment of PIM1 kinase activity (Supplemental Figure 7), we demonstrated a 7-fold increase in Pim-1 from cCICs-Pim-1 versus control cCICs (Figure 6). We then used a panel of sensors designed for other kinases relevant to PIM-1 signaling and observed increases in lysates from cCICs versus control cCICs for CK1 (1.8-fold), IKK-family (2.7-fold), AKT/SGK (3.1-fold), CDK1-3/5 (3.7-fold), p38/MAPKs (2.8-fold) and ERK1/2 MAPKs (2.6-fold) (Figure 7 and Table 2). In contrast, we saw no increase in activity with a sensor that measures the activity of JNK1, 2 or 3 (Figure 7 and Table 2). These changes are consistent with heightened kinase activity resulting from Pim-1 overexpression, reported by the Sussman laboratory and others, that contributes to Pim-1-mediated effects. Importantly, these changes are all strikingly lower than the observed increase in PIM-1 activity, which presumably highlights an important balance in post-transcriptional and post-translational influences established in these cCICs that may help explain the beneficial effect of Pim-1 genetic engineering to enhance cell survival and proliferation, while avoiding any negative consequences, such as an oncogenesis. This conclusion is consistent with those previously proposed and reported by the Sussman laboratory.^20^

**Table 2.** PIM-1 kinase pathway activity

| Crude Cell Lysate Sample | Sensor (10 μM) | Target Kinase(s) | Reaction Rate | | Fold Change |
| --- | --- | --- | --- | --- | --- |
| | | | RFU/min/μg | Std Dev | |
| HI3067/P12/GFP | AQT0001 | Tyr Kinases | 4.3 | 1.1 | 0.6 |
| HI3067/P12/PIM1 | AQT0001 | Tyr Kinases | 2.78 | 0.40 |  |
| HI3067/P12/GFP | ST24 | CK1 | 15.2 | 1 | 1.8 |
| HI3067/P12/PIM1 | ST24 | CK1 | 26.90 | 3.80 |  |
| HI3067/P12/GFP | AQT215 | IKK/TBK | 5.2 | 1.5 | 2.7 |
| HI3067/P12/PIM1 | AQT215 | IKK/TBK | 13.80 | 1.60 |  |
| HI3067/P12/GFP | AQT0233 | AKT/SGK | 9.0 | 2.7 | 3.1 |
| HI3067/P12/PIM1 | AQT0233 | AKT/SGK | 28.27 | 3.90 |  |
| HI3067/P12/GFP | AQT0255 | CDK1-3,5 | 16.6 | 1.7 | 3.7 |
| HI3067/P12/PIM1 | AQT0255 | CDK1-3,5 | 60.82 | 3.50 |  |
| HI3067/P12/GFP | AQT0258 | CDK4,6 | 13.3 | 3.8 | 1.0 |
| HI3067/P12/PIM1 | AQT0258 | CDK4,6 | 13.70 | 5.40 |  |
| HI3067/P12/GFP | AQT0365 | JNK1-3 | 5.7 | 2.7 | 1.2 |
| HI3067/P12/PIM1 | AQT0365 | JNK1-3 | 6.96 | 0.70 |  |
| HI3067/P12/GFP | AQT0374 | p38 | 44.0 | 3.2 | 2.8 |
| HI3067/P12/PIM1 | AQT0374 | p38 | 124.77 | 8.30 |  |
| HI3067/P12/GFP | AQT0376 | ERK1/2 | 83.8 | 4.3 | 2.6 |
| HI3067/P12/PIM1 | AQT0376 | ERK1/2 | 221.20 | 28.50 |  |
| HI3067/P12/GFP | AQT57218 1C09 | PIM1 (M3) | 6.0 | 1 | 7.4 |
| HI3067/P12/PIM1 | AQT57218 1C09 | PIM1 (M3) | 44.16 | 3.10 |  |

**Figure 6.**
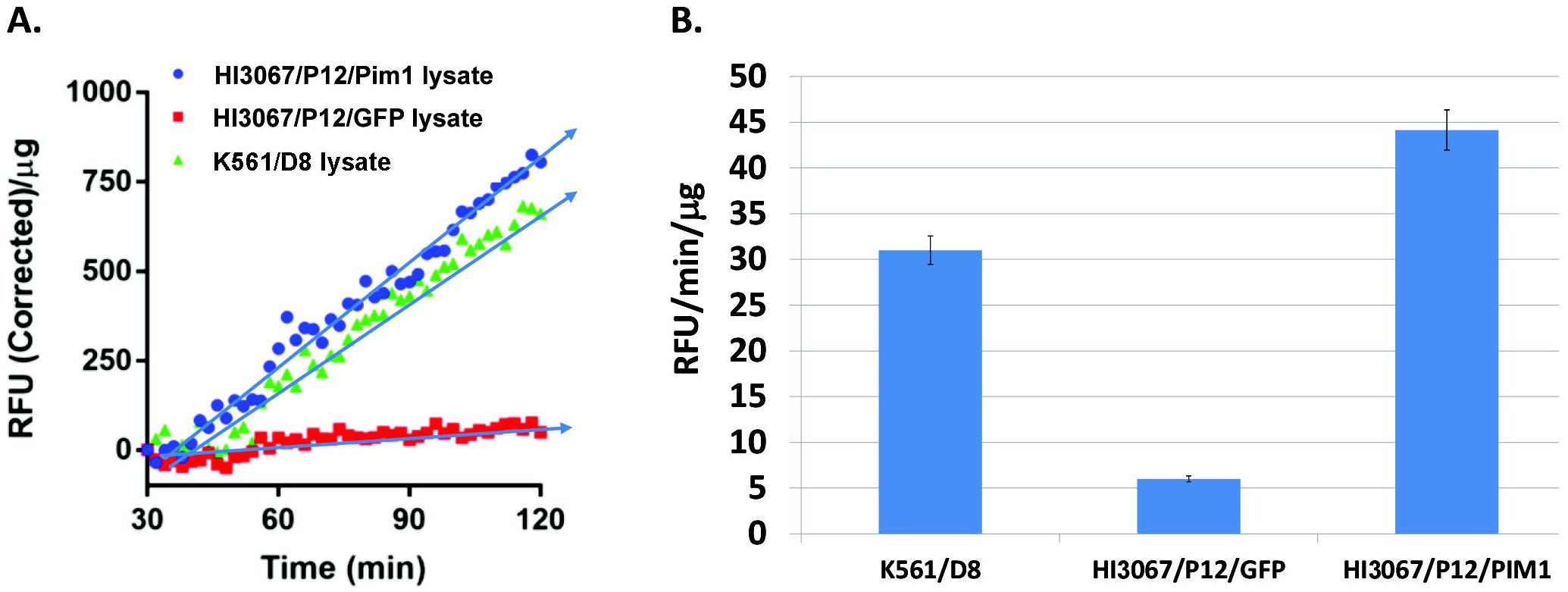
PIM-1 kinase pathway activity elevation resulting from Pim-1 overexpression. (A) Kinase reactions were monitored kinetically and the net fluorescence signal (RFU Corrected/μg of crude cell lysate protein) determined and plotted versus time. The slope of the linear region for each curve between 30 and 120 minutes was determined using GraphPad Prism software and (B) measure of the initial reaction rate (RFU Corrected /min/μg of crude cell lysate protein).

**Figure 7.**
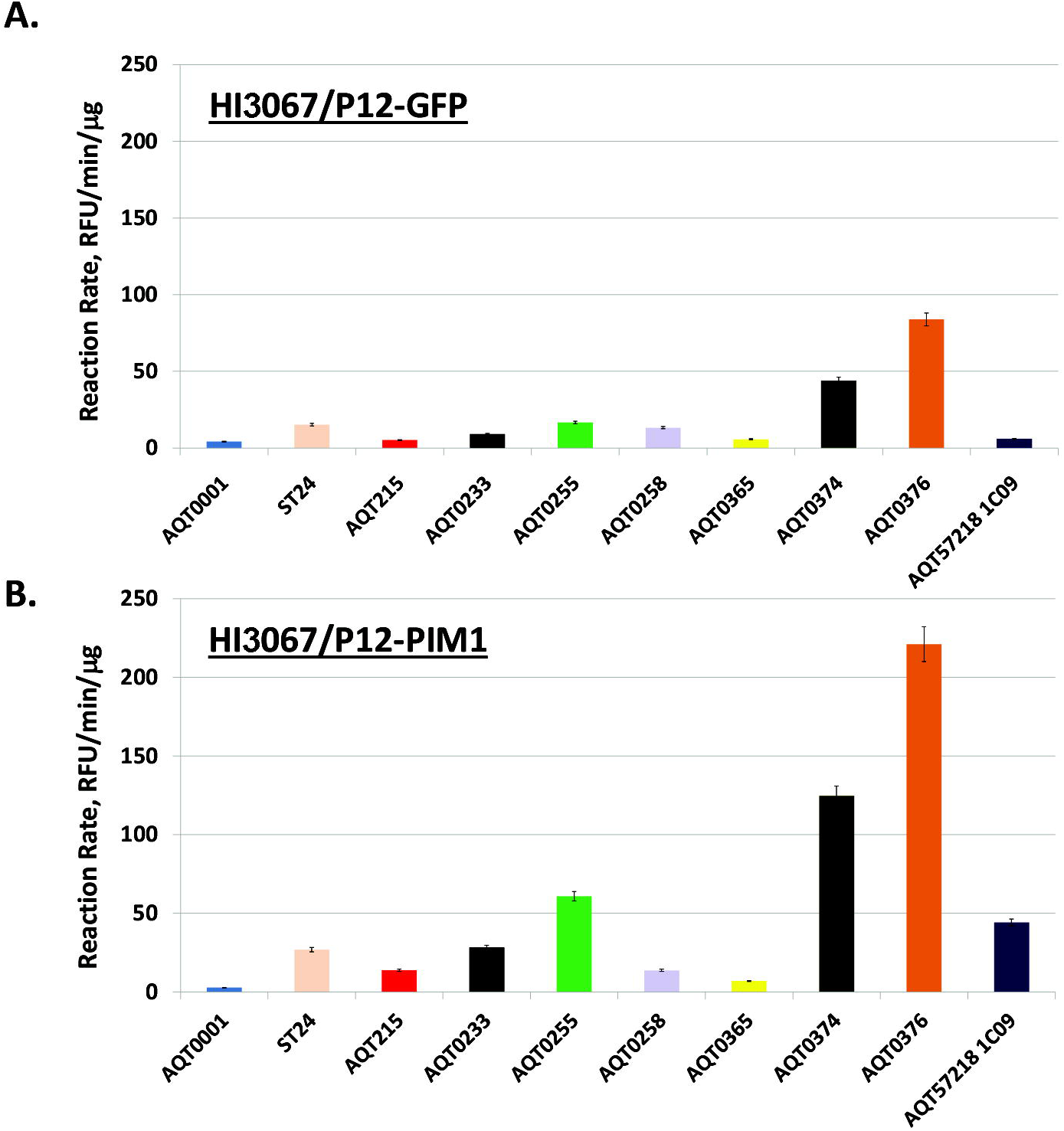
Kinase pathway activity elevated from Pim-1 overexpression. Kinase reactions were monitored kinetically and the initial reaction rate (RFU Corrected /min/μg of crude cell lysate protein) was determined for each sensor in (A) control cCICs and (B) cCICs-Pim-1. Highest activation is seen with AQT0215 (IKKs), AQT0233 (AKT/SGK), AQT0255 (CDK1-3), AQT0374 (p38), AQT0376 (ERK1/2), and AQT57218-1C09 (PIM1 & 3).

## Discussion

Next generation stem cell therapeutic approaches will require innovative solutions to current limitations in implementation. A universally accepted and serious challenge for efficacy is poor survival, engraftment, and persistence of adoptively transferred cells, which is poor in the setting of autologous treatment and arguably even worse in allogeneic therapy. Toward the goal of confronting these longstanding issues, our group embarked upon a molecular interventional strategy to enhance stem cell therapy through genetic engineering with pro-survival and pro-proliferative Pim-1 kinase. Myocardial Pim-1 biology as reported since 2007 spawned 64 peer review articles including 24 publications from the Sussman laboratory (PubMed key word search for “Pim-1” and “cardiac”). As such, the mechanism of Pim-1 action is likely documented equal to or better than any other cardiac regenerative therapy. Pim-1 expression is stimulated by a variety of hormones, cytokines and mitogens, many of which are associated with cardioprotective signaling [38,39]. Despite pro-proliferative effects mediated by Pim-1, oncogenic transformation has never been observed in any of our samples, and all Pim-1 enhanced cCICs were amenable to differentiation *in vitro* that resulted in acquisition of post-mitotic characteristics when injected into rodents or swine. Histological evaluation findings for oncogenic transformation, tumorigenicity, teratomas, ectopic tissue formation or similar neoplastic plastic processes were all negative for all cCICs treated animals on whole body necropsy either at 4- or 8-weeks following study product administration.^22^

Human cCICs overexpressing Pim-1 show no evidence of oncogenicity in NOD/SCID mice^23^. Data from this validation study revealed trafficking, distribution, and persistence of cCICs-Pim-1 for up to 32 weeks. Negligible risk from Pim-1 modification is further supported by the following findings: A) Pim-1 literature inhibiting activity in cancer reinforces Pim-1 function as a cell survival enhancer after undergoing an oncogenic event rather than an initiator of transformation. Likelihood for oncogenic transformation is very remote because of cell-type effects and context-dependent action. PIM-1 is a weak oncogene at best because it is not a transforming factor by itself as supported by published literature.^41,68-70^ B) High level Pim-1 overexpression in cCICs is cytotoxic rather than oncogenic and we are unable to expand cardiac interstitial cells exhibiting high Pim-1 overexpression. C) The CMV promoter employed in the lentiviral vector is progressively down-regulated *in vivo*, likely by epigenetic modification (e.g. methylation) such that Pim-1 levels in cCICs-Pim-1 is increasingly difficult to discriminate from endogenous myocardium within weeks after adoptive transfer.^22^ D) cCICs-Pim-1 undergo replicative senescence near 20 passages in culture and cannot be further expanded.^25^ E) c-Myc levels are low in cardiac stem cells and unchanged in cCICs-Pim-1 reinforcing their low oncogenic potential. F) Clinical trials use embryonic stem cells or induced pluripotent stem cells which possess demonstrable oncogenic potential. In contrast, adult stem cells used in clinical therapy for decades have an outstanding safety record, highlighting low oncogenic potential of adult stem cells. G) The heart failure target population consists of primarily elderly individuals with age-limited stem cell proliferative and survival capacity, highlighting the rationale for cCICs-Pim-1 therapy. H) Concerns regarding potential for cardiac adult stem cells to “transdifferentiate” into another stem cell type of distinct origin has no precedent in current literature or our laboratory findings. Cardiac stem cell biology is vastly different from hematopoietic stem cells and collective observations indicate cardiac versus hematopoietic stem cells are very distinct lineages. Specifically, transcriptional profiling is distinct^71,72^, cardiac stem cells die in hematopoietic stem cell culture media and vice-versa, and hematopoietic stem cells die in the myocardium quickly within days of arrival whereas cardiac stem cells persist. Pim-1 overexpression impact is cell-type specific and after modifying innumerable cardiac stem cells with Pim-1 over many years Pim-1 a transformed phenotype has not arisen. I) No evidence exists that non-engrafted cCICs-Pim-1 lodge in alternative non-cardiac sites despite years of research injecting the cells into both NOD-SCID mice as well as immunosuppressed miniswine. J) Karyotype analysis demonstrated human cCICs-Pim-1 to be normal without chromosomal aberrations.^23^ Multiple potential explanations likely account for lack of a metastatic phenotype for cCICs-Pim-1 that stands in contrast to unrelated observations pertinent to embryonic stem cells or iPS cells. Specifically, augmented regenerative potential means that cCICs-Pim-1 are delivered at lower cell dosing relative to alternative cell types, and cCICs-Pim-1 are predisposed to cardiogenic commitment as evident by transcriptional profiling as well as protein expression and therefore likely undergo apoptotic death by anoikis if lodged out of context in non-myocardial tissue. Nevertheless, despite this collective body of evidence concerns continue to be articulated that oncogenic risk resulting from Pim-1 overexpression is a potential issue.^30^

There is no clear consensus for an optimal experimental model to definitely assess vector safety for a particular cell engineering application. Lentiviral vectors have integration sites away from transcriptional regulatory sites and low oncogenic risk and to quote: “…tumorigenesis was unaffected by lentiviral vectors, despite a higher integration load and robust expression of lentiviral vectors in all hematopoietic lineages. The results of these studies have been confirmed by others and indicate a favorable safety profile for lentiviral vectors. Current clinical experience with lentiviral vectors supports these experimental studies. There have been no oncogenic events observed in human clinical trials using lentiviral vectors to date.”^73^. Regarding lentiviral oncogenic risk, consider that lentiviral vectors have made their way into clinics as therapies, including for advanced forms of HIV infections^74^, Parkinson’s disease^75^, and inherited disorders affecting hematopoietic cells.^76^ In addition, lentiviral vectors have integration sites away from transcriptional regulatory sites, making them a safe therapeutic option^33^. These findings are in stark contrast to published literature showing chromosomal abnormalities in certain embryonic stem cells and induced pluripotent stem cells where oncogenesis remains a significant barrier to therapeutic implementation.^77^ Prior studies with hematopoietic stem cells that, unlike cCICs, normally undergo complex genetic rearrangements as part of their differentiation program exhibited tumorigenic potential from induction of cellular oncogenes adjacent to the integration site^33^.

Heart failure associated with aging has been proposed to be a “stem cell disease” characterized by impaired functional reserve of the endogenous stem cell pool due to exhaustion, senescence, depletion, or inability to cope with the environmental stressors.^78^ Stem cell therapy for cardiac repair holds great promise, but the ability of cCICs to repair damaged myocardium declines with age.^78-80^ Shortening of telomere length has been linked to both senescence^81^ and cell death^82^, further highlighting concerns related to aging and pathological stress. cCICs derived from patients with advanced biological age and severe concurrent clinical features will require rejuvenation to reverse deleterious effects of aging and disease. Lack of overt risk of oncogenic transformation as well as the *ex vivo* approach to genetic modification described in this study that allows for rigorous screening prior to reintroduction into a patient for autologous therapy provides reassurance of safety for use of Pim-1 mediated enhancement for cCICs therapeutic use.

## Acknowledgements

The research in this study was funded by NIH STTR R41HL1377602 to CardioCreate Inc. We thank the staff at Sharp Hospital, in particular Chris Kohlmeyer and Donna Small, for bridging the collaborative arrangement between Sharp Hospital and the Sussman Laboratory. K.M. Broughton is supported by NIH grant F32HL136196. M.A. Sussman is also supported by NIH grants: R01HL067245, R37HL091102, R01HL105759, R01HL113647, R01HL117163, P01HL085577, and R01HL122525, as well as an award from the Fondation Leducq.

## References

1 Chen, C., Termglinchan, V. & Karakikes, I. Concise Review: Mending a Broken Heart: The Evolution of Biological Therapeutics. Stem Cells 35, 1131–1140, doi:10.1002/stem.2602 (2017).

2 Crowley, M. G. & Tajiri, N. Exogenous stem cells pioneer a biobridge to the advantage of host brain cells following stroke: New insights for clinical applications. Brain Circ 3, 130–134, doi:10.4103/bc.bc_17_17 (2017).

3 Goldberg, A., Mitchell, K., Soans, J., Kim, L. & Zaidi, R. The use of mesenchymal stem cells for cartilage repair and regeneration: a systematic review. J Orthop Surg Res 12, 39, doi:10.1186/s13018-017-0534-y (2017).

4 Jantzie, L. L., Scafidi, J. & Robinson, S. Stem cells and cell-based therapies for cerebral palsy: a call for rigor. Pediatr Res 83, 345–355, doi:10.1038/pr.2017.233 (2018).

5 Krueger, T. E. G., Thorek, D. L. J., Denmeade, S. R., Isaacs, J. T. & Brennen, W. N. Concise Review: Mesenchymal Stem Cell-Based Drug Delivery: The Good, the Bad, the Ugly, and the Promise. Stem Cells Transl Med 7, 651–663, doi:10.1002/sctm.18-0024 (2018).

6 Quesenberry, P. & Goldberg, L. R. A New Stem Cell Biology: Transplantation and Baseline, Cell Cycle and Exosomes. Adv Exp Med Biol 1056, 3–9, doi:10.1007/978-3-319- 74470-4_1 (2018).

7 Vinhas, A., Rodrigues, M. T. & Gomes, M. E. Exploring Stem Cells and Inflammation in Tendon Repair and Regeneration. Adv Exp Med Biol 1089, 37–46, doi:10.1007/5584_2018_258 (2018).

8 Witman, N. & Sahara, M. Cardiac Progenitor Cells in Basic Biology and Regenerative Medicine. Stem Cells Int 2018, 8283648, doi:10.1155/2018/8283648 (2018).

9 Youssef, A. A. et al. The Promise and Challenge of Induced Pluripotent Stem Cells for Cardiovascular Applications. JACC Basic Transl Sci 1, 510–523, doi:10.1016/j.jacbts.2016.06.010 (2016).

10 Zhu, Y., Chen, X., Yang, X. & Ei-Hashash, A. Stem cells in lung repair and regeneration: Current applications and future promise. J Cell Physiol 233, 6414–6424, doi:10.1002/jcp.26414 (2018).

11 Aagaard, K. S., Ganesalingam, S., Jensen, C. H., Sheikh, S. P. & Andersen, D. C. Poor engraftment potential of epicardial progenitors upon intramyocardial transplantation into the neonatal mouse heart. Int J Cardiol 168, 4360–4362, doi:10.1016/j.ijcard.2013.05.061 (2013).

12 Der Sarkissian, S., Levesque, T. & Noiseux, N. Optimizing stem cells for cardiac repair: Current status and new frontiers in regenerative cardiology. World J Stem Cells 9, 9–25, doi:10.4252/wjsc.v9.i1.9 (2017).

13 Hong, K. U. et al. c-kit+ Cardiac stem cells alleviate post-myocardial infarction left ventricular dysfunction despite poor engraftment and negligible retention in the recipient heart. PLoS One 9, e96725, doi:10.1371/journal.pone.0096725 (2014).

14 Kanda, P. & Davis, D. R. Cellular mechanisms underlying cardiac engraftment of stem cells. Expert Opin Biol Ther 17, 1127–1143, doi:10.1080/14712598.2017.1346080 (2017).

15 Li, X., Tamama, K., Xie, X. & Guan, J. Improving Cell Engraftment in Cardiac Stem Cell Therapy. Stem Cells Int 2016, 7168797, doi:10.1155/2016/7168797 (2016).

16 Serpooshan, V. & Wu, S. M. Patching up broken hearts: cardiac cell therapy gets a bioengineered boost. Cell Stem Cell 15, 671–673, doi:10.1016/j.stem.2014.11.008 (2014).

17 Yanamandala, M. et al. Overcoming the Roadblocks to Cardiac Cell Therapy Using Tissue Engineering. J Am Coll Cardiol 70, 766–775, doi:10.1016/j.jacc.2017.06.012 (2017).

18 Mohsin, S., Siddiqi, S., Collins, B. & Sussman, M. A. Empowering adult stem cells for myocardial regeneration. Circ Res 109, 1415–1428, doi:10.1161/CIRCRESAHA.111.243071 (2011).

19 Muraski, J. A. et al. Pim-1 regulates cardiomyocyte survival downstream of Akt. Nat Med 13, 1467–1475, doi:10.1038/nm1671 (2007).

20 Siddiqi, S. & Sussman, M. A. Cell and gene therapy for severe heart failure patients: the time and place for Pim-1 kinase. Expert Rev Cardiovasc Ther 11, 949–957, doi:10.1586/14779072.2013.814830 (2013).

21 Cottage, C. T. et al. Cardiac progenitor cell cycling stimulated by pim-1 kinase. Circ Res 106, 891–901, doi:10.1161/CIRCRESAHA.109.208629 (2010).

22 Kulandavelu, S. et al. Pim1 Kinase Overexpression Enhances ckit(+) Cardiac Stem Cell Cardiac Repair Following Myocardial Infarction in Swine. J Am Coll Cardiol 68, 2454–2464, doi:10.1016/j.jacc.2016.09.925 (2016).

23 Mohsin, S. et al. Human cardiac progenitor cells engineered with Pim-I kinase enhance myocardial repair. J Am Coll Cardiol 60, 1278–1287, doi:10.1016/j.jacc.2012.04.047 (2012).

24 Fischer, K. M. et al. Pim-1 kinase inhibits pathological injury by promoting cardioprotective signaling. J Mol Cell Cardiol 51, 554–558, doi:10.1016/j.yjmcc.2011.01.004 (2011).

25 Mohsin, S. et al. Rejuvenation of human cardiac progenitor cells with Pim-1 kinase. Circ Res 113, 1169–1179, doi:10.1161/CIRCRESAHA.113.302302 (2013).

26 Broughton, K. M. & Sussman, M. A. Enhancement Strategies for Cardiac Regenerative Cell Therapy: Focus on Adult Stem Cells. Circ Res 123, 177–187, doi:10.1161/CIRCRESAHA.118.311207 (2018).

27 Brunen, D., de Vries, R. C., Lieftink, C., Beijersbergen, R. L. & Bernards, R. PIM Kinases Are a Potential Prognostic Biomarker and Therapeutic Target in Neuroblastoma. Mol Cancer Ther 17, 849–857, doi:10.1158/1535-7163.MCT-17-0868 (2018).

28 Jimenez-Garcia, M. P. et al. The role of PIM1/PIM2 kinases in tumors of the male reproductive system. Sci Rep 6, 38079, doi:10.1038/srep38079 (2016).

29 Mondello, P., Cuzzocrea, S. & Mian, M. Pim kinases in hematological malignancies: where are we now and where are we going? J Hematol Oncol 7, 95, doi:10.1186/s13045-014- 0095-z (2014).

30 Hinkel, R. Pim1 Overexpressing ckit(+) Cardiac Stem Cells in Cardiac Regeneration: Preconditioning as Next-Generation Stem Cell Therapy? J Am Coll Cardiol 68, 2465–2466, doi:10.1016/j.jacc.2016.09.924 (2016).

31 Bokhoven, M. et al. Insertional gene activation by lentiviral and gammaretroviral vectors. J Virol 83, 283–294, doi:10.1128/JVI.01865-08 (2009).

32 Cesana, D. et al. Uncovering and dissecting the genotoxicity of self-inactivating lentiviral vectors in vivo. Mol Ther 22, 774–785, doi:10.1038/mt.2014.3 (2014).

33 Modlich, U. et al. Insertional transformation of hematopoietic cells by self-inactivating lentiviral and gammaretroviral vectors. Mol Ther 17, 1919–1928, doi:10.1038/mt.2009.179 (2009).

34 Montini, E. et al. Hematopoietic stem cell gene transfer in a tumor-prone mouse model uncovers low genotoxicity of lentiviral vector integration. Nat Biotechnol 24, 687–696, doi:10.1038/nbt1216 (2006).

35 Ranzani, M. et al. Lentiviral vector-based insertional mutagenesis identifies genes associated with liver cancer. Nat Methods 10, 155–161, doi:10.1038/nmeth.2331 (2013).

36 Wang, Z. et al. Pim-1: a serine/threonine kinase with a role in cell survival, proliferation, differentiation and tumorigenesis. J Vet Sci 2, 167–179 (2001).

37 Domen, J. et al. Comparison of the human and mouse PIM-1 cDNAs: nucleotide sequence and immunological identification of the in vitro synthesized PIM-1 protein. Oncogene Res 1, 103–112 (1987).

38 van Lohuizen, M. et al. Predisposition to lymphomagenesis in pim-1 transgenic mice: cooperation with c-myc and N-myc in murine leukemia virus-induced tumors. Cell 56, 673–682 (1989).

39 Baron, B. W. et al. PIM1 gene cooperates with human BCL6 gene to promote the development of lymphomas. Proc Natl Acad Sci U S A 109, 5735–5739, doi:10.1073/pnas.1201168109 (2012).

40 Moroy, T., Grzeschiczek, A., Petzold, S. & Hartmann, K. U. Expression of a Pim-1 transgene accelerates lymphoproliferation and inhibits apoptosis in lpr/lpr mice. Proc Natl Acad Sci U S A 90, 10734–10738 (1993).

41 Narlik-Grassow, M. et al. Conditional transgenic expression of PIM1 kinase in prostate induces inflammation-dependent neoplasia. PLoS One 8, e60277, doi:10.1371/journal.pone.0060277 (2013).

42 Bhattacharya, N. et al. Pim-1 associates with protein complexes necessary for mitosis. Chromosoma 111, 80–95, doi:10.1007/s00412-002-0192-6 (2002).

43 Wang, Z. et al. Phosphorylation of the cell cycle inhibitor p21Cip1/WAF1 by Pim-1 kinase. Biochim Biophys Acta 1593, 45–55 (2002).

44 Weirauch, U. et al. Functional role and therapeutic potential of the pim-1 kinase in colon carcinoma. Neoplasia 15, 783–794 (2013).

45 Zhang, Y., Wang, Z., Li, X. & Magnuson, N. S. Pim kinase-dependent inhibition of c-Myc degradation. Oncogene 27, 4809–4819, doi:10.1038/onc.2008.123 (2008).

46 Lilly, M., Sandholm, J., Cooper, J. J., Koskinen, P. J. & Kraft, A. The PIM-1 serine kinase prolongs survival and inhibits apoptosis-related mitochondrial dysfunction in part through a bcl-2-dependent pathway. Oncogene 18, 4022–4031, doi:10.1038/sj.onc.1202741 (1999).

47 Gu, J. J., Wang, Z., Reeves, R. & Magnuson, N. S. PIM1 phosphorylates and negatively regulates ASK1-mediated apoptosis. Oncogene 28, 4261–4271, doi:10.1038/onc.2009.276 (2009).

48 Kim, K. T. et al. Pim-1 is up-regulated by constitutively activated FLT3 and plays a role in FLT3-mediated cell survival. Blood 105, 1759–1767, doi:10.1182/blood-2004-05-2006 (2005).

49 White, E. The pims and outs of survival signaling: role for the Pim-2 protein kinase in the suppression of apoptosis by cytokines. Genes Dev 17, 1813–1816, doi:10.1101/gad.1123103 (2003).

50 Monsanto, M. M. et al. Concurrent Isolation of 3 Distinct Cardiac Stem Cell Populations From a Single Human Heart Biopsy. Circ Res 121, 113–124, doi:10.1161/CIRCRESAHA.116.310494 (2017).

51 Pertea, M., Kim, D., Pertea, G. M., Leek, J. T. & Salzberg, S. L. Transcript-level expression analysis of RNA-seq experiments with HISAT, StringTie and Ballgown. Nat Protoc 11, 1650–1667, doi:10.1038/nprot.2016.095 (2016).

52 Lukovic, E., Gonzalez-Vera, J. A. & Imperiali, B. Recognition-domain focused chemosensors: versatile and efficient reporters of protein kinase activity. J Am Chem Soc 130, 12821–12827, doi:10.1021/ja8046188 (2008).

53 Lukovic, E., Vogel Taylor, E. & Imperiali, B. Monitoring protein kinases in cellular media with highly selective chimeric reporters. Angew Chem Int Ed Engl 48, 6828–6831, doi:10.1002/anie.200902374 (2009).

54 Peterson, L. B., Yaffe, M. B. & Imperiali, B. Selective mitogen activated protein kinase activity sensors through the application of directionally programmable D domain motifs. Biochemistry 53, 5771–5778, doi:10.1021/bi500862c (2014).

55 Shults, M. D., Carrico-Moniz, D. & Imperiali, B. Optimal Sox-based fluorescent chemosensor design for serine/threonine protein kinases. Anal Biochem 352, 198–207, doi:10.1016/j.ab.2006.03.003 (2006).

56 Shults, M. D. & Imperiali, B. Versatile fluorescence probes of protein kinase activity. J Am Chem Soc 125, 14248–14249, doi:10.1021/ja0380502 (2003).

57 Shults, M. D., Janes, K. A., Lauffenburger, D. A. & Imperiali, B. A multiplexed homogeneous fluorescence-based assay for protein kinase activity in cell lysates. Nat Methods 2, 277–283, doi:10.1038/nmeth747 (2005).

58 Stains, C. I. et al. Interrogating signaling nodes involved in cellular transformations using kinase activity probes. Chem Biol 19, 210–217, doi:10.1016/j.chembiol.2011.11.012 (2012).

59 Borowicz, S. et al. The soft agar colony formation assay. J Vis Exp, e51998, doi:10.3791/51998 (2014).

60 Desgrosellier, J. S. et al. An integrin alpha(v)beta(3)-c-Src oncogenic unit promotes anchorage-independence and tumor progression. Nat Med 15, 1163–1169, doi:10.1038/nm.2009 (2009).

61 Bonassi, S. et al. HUman MicroNucleus project: international database comparison for results with the cytokinesis-block micronucleus assay in human lymphocytes: I. Effect of laboratory protocol, scoring criteria, and host factors on the frequency of micronuclei. Environ Mol Mutagen 37, 31–45 (2001).

62 Fenech, M. Cytokinesis-block micronucleus cytome assay. Nat Protoc 2, 1084–1104, doi:10.1038/nprot.2007.77 (2007).

63 Fenech, M., Holland, N., Chang, W. P., Zeiger, E. & Bonassi, S. The HUman MicroNucleus Project-An international collaborative study on the use of the micronucleus technique for measuring DNA damage in humans. Mutat Res 428, 271–283 (1999).

64 Kirsch-Volders, M. et al. In vitro genotoxicity testing using the micronucleus assay in cell lines, human lymphocytes and 3D human skin models. Mutagenesis 26, 177–184, doi:10.1093/mutage/geq068 (2011).

65 Le Hegarat, L. et al. Assessment of the genotoxic potential of indirect chemical mutagens in HepaRG cells by the comet and the cytokinesis-block micronucleus assays. Mutagenesis 25, 555–560, doi:10.1093/mutage/geq039 (2010).

66 Le Hegarat, L. et al. Performance of comet and micronucleus assays in metabolic competent HepaRG cells to predict in vivo genotoxicity. Toxicol Sci 138, 300–309, doi:10.1093/toxsci/kfu004 (2014).

67 Stains, C. I., Lukovic, E. & Imperiali, B. A p38alpha-selective chemosensor for use in unfractionated cell lysates. ACS Chem Biol 6, 101–105, doi:10.1021/cb100230y (2011).

68 Biffi, A. et al. Lentiviral vector common integration sites in preclinical models and a clinical trial reflect a benign integration bias and not oncogenic selection. Blood 117, 5332–5339, doi:10.1182/blood-2010-09-306761 (2011).

69 Aguirre, E., Renner, O., Narlik-Grassow, M. & Blanco-Aparicio, C. Genetic Modeling of PIM Proteins in Cancer: Proviral Tagging and Cooperation with Oncogenes, Tumor Suppressor Genes, and Carcinogens. Front Oncol 4, 109, doi:10.3389/fonc.2014.00109 (2014).

70 Holder, S. L. & Abdulkadir, S. A. PIM1 kinase as a target in prostate cancer: roles in tumorigenesis, castration resistance, and docetaxel resistance. Curr Cancer Drug Targets 14, 105–114 (2014).

71 Hammerman, P. S., Fox, C. J., Birnbaum, M. J. & Thompson, C. B. Pim and Akt oncogenes are independent regulators of hematopoietic cell growth and survival. Blood 105, 4477–4483, doi:10.1182/blood-2004-09-3706 (2005).

72 Rossini, A. et al. Human cardiac and bone marrow stromal cells exhibit distinctive properties related to their origin. Cardiovasc Res 89, 650–660, doi:10.1093/cvr/cvq290 (2011).

73 Leath, A. & Cornetta, K. Developing novel lentiviral vectors into clinical products. Methods Enzymol 507, 89–108, doi:10.1016/B978-0-12-386509-0.00005-3 (2012).

74 Mautino, M. R. Lentiviral vectors for gene therapy of HIV-1 infection. Curr Gene Ther 2, 23–43 (2002).

75 Lundberg, C. et al. Applications of lentiviral vectors for biology and gene therapy of neurological disorders. Curr Gene Ther 8, 461–473 (2008).

76 Woods, N. B., Ooka, A. & Karlsson, S. Development of gene therapy for hematopoietic stem cells using lentiviral vectors. Leukemia 16, 563–569, doi:10.1038/sj.leu.2402447 (2002).

77 Ahmed, R. P., Ashraf, M., Buccini, S., Shujia, J. & Haider, H. Cardiac tumorigenic potential of induced pluripotent stem cells in an immunocompetent host with myocardial infarction. Regen Med 6, 171–178, doi:10.2217/rme.10.103 (2011).

78 Cesselli, D. et al. Effects of age and heart failure on human cardiac stem cell function. Am J Pathol 179, 349–366, doi:10.1016/j.ajpath.2011.03.036 (2011).

79 Capogrossi, M. C. Cardiac stem cells fail with aging: a new mechanism for the age-dependent decline in cardiac function. Circ Res 94, 411–413, doi:10.1161/01.RES.0000122070.37999.1B (2004).

80 Dimmeler, S. & Leri, A. Aging and disease as modifiers of efficacy of cell therapy. Circ Res 102, 1319–1330, doi:10.1161/CIRCRESAHA.108.175943 (2008).

81 Fyhrquist, F., Saijonmaa, O. & Strandberg, T. The roles of senescence and telomere shortening in cardiovascular disease. Nat Rev Cardiol 10, 274–283, doi:10.1038/nrcardio.2013.30 (2013).

82 Booth, S. A. & Charchar, F. J. Cardiac telomere length in heart development, function, and disease. Physiol Genomics 49, 368–384, doi:10.1152/physiolgenomics.00024.2017 (2017).

