## Supplement for "Safety profiling of genetically engineered Pim-1 kinase overexpression for oncogenicity risk in human c-kit+ cardiac interstitial cells"

### Supplemental Methods

**Lentiviral Vector Production Protocol* Adjusted from *Production of CGMP-Grade Lentiviral Vectors*, Lara J. Ausubel, et al. 2012**

**Reagents:**

1. Cells HEK 293T/17 (ATCC Item number CRL-11268; Batch number 63696280). Certificate of analysis included upon request. Hereon, referred to as HEK 293T's.
2. Packaging plasmids pantropic ViraSafe™ 3rd generation lentiviral packaging system (Cell Biolabs, INC catalog number VPK-206). Plasmid maps included upon request. Components: pRSV-Rev Packaging Vector (referred to as Rev) Part number 320022, pCMV-VSV-G Envelope Vector (referred to as VSVG) Part number RV-110, pCgpV Packaging Vector (referred to as g/p) Part number 320024.
3. 3rd generation transfer vectors: 1. CGW Pim1 eGFP, Pim1 under CMV regulation, GFP downstream of IRES. 2. Control vector CGW eGFP downstream of IRES. Plasmid maps available upon request.
4. Cell culture medium for HEK 293T's with all components being cGMP compliant. Ingredients: DMEM with high glucose (4.5g/L), 1% sodium pyruvate and L-Glutamine (Gibco catalog number 11995040) supplemented with 10% fetal bovine serum (Omega Scientific catalog number FB-01; lot# 703107), penicillin-streptomycin 100 units and 100μg per mL of medium respectively (Lonza catalog number 17-602F; lot number 16J115303). Referred to as base medium
5. 1:1 trypsin:cell stripper.
6. PBS (phosphate buffered saline), pH 7.4 (Gibco catalog number 10010023)
7. PEI polyethylenimine "Max", (Mw 40,000)- high potency linear (polysciences catalog number 24765-1) 1mg/mL dissolved in cell culture grade water (Hyclone from Fisher Scientific catalog number SH30529.LS), pH adjusted by 1N HCL, followed by sterile filtration (0.22 μm stericup, Millipore catalog number, SCGPU05RE). Aliquot and store at -20°C.
8. Benzonase endonuclease (Millipore catalog number 1.01654.0001)
9. Magnesium Chloride solution, 1M (Millipore catalog number 5985-OP)

*Following CGMP regulations in a research laboratory setting.

**Plasticware:**

10. 10-stack (6360 $cm^2$) Corning CELLSTACK Culture Chambers (Corning, catalog number 3270, Fisher Scientific, catalog number 05-539-096)
11. Olympus T75 (Genesee Scientific, catalog number 25-209 and T182 vented flasks (Genesee Scientific, catalog number 25-211)
12. Corning disposable sterile bottles, 10 L (Fisher Scientific catalog number 09-761-11), 150 mL (Fisher Scientific catalog number 09-761-140)
13. Sterile 50 mL conical centrifuge tubes (Genesee Scientific, catalog number 28-106)
14. Sterile micro tube 0.5 mL with attached screwed cap (Sarstedt, catalog number 72.730.106)
15. 500 mL Stericup filter 0.22 $\mu$L (Millipore, catalog number SCGPU05RE), 250 mL Stericup filter 0.22 $\mu$L (Millipore, catalog number SCGPU02RE), 250 mL Stericup filter 0.45 $\mu$L (Millipore, catalog number SCHVU02RE)
16. Thinwall polypropylene conical tubes (Beckman Coulter, catalog number 358126)

**Lentiviral Particle Production Protocol:**

**All steps are conducted under laminar flow following sterile method.**

**Day 1:** Prior to day 1 expand HEK 293T's to yield ~2 x $10^8$ cells on day 1. Wash the cells with 1 x PBS, dislodge and pellet by centrifugation. Count the cells and plate 3 x $10^4$ cells/$cm^2$ in 100$\mu$L/$cm^2$ base medium on a 10-stack cell culture (6,360 $cm^2$) flask (Coring). Total medium volume 636 mL, total cell number 1.9 x $10^8$. Include a T75 as a mock plate, with 30 x $10^4$ cells per $cm^2$ and 100$\mu$L/$cm^2$ cell culture base medium.

**Day 3:** In the evening ~04:00-05:00PM start making the transfection mixture. Add all of the packaging plasmids and transfer plasmid (for volumes refer to Supplemental Table 2**)** into 150 mL sterile bottle. Add 105.8 mL of 1 x DMEM (room temperature) with no additives. Gently homogenize the solution using a 50 mL serological pipette. Equally divide the contents into 3 x 50 mL tubes (total volume of plasmid mixture 109.7 mL divided by 3 equals 36.6 mL) Vortex and centrifuge for 1 minute 1,200 rpm. Prepare a 50 mL conical tube with 21.2 mL of 1 x DMEM with no additives and add 5.3 mL of PEI (1mg/mL) directly into the medium, vortex and centrifuge for 1 minute 1,200 rpm. Incubate the 150 mL and 50 mL tubes for 5 minutes at room temperature. Add 8.8 mL PEI mixture to the plasmid tubes. Vortex the tube for 10 seconds following a 1 minute 1,200 rpm centrifugation and a 15-minute incubation at room temperature. Total volume ~ 136.2 mL. Add 1.6 mL of plasmid mixture into 5.9 mL of 37°C base medium for the mock plate. The rest of the plasmid and PEI mixture add to 501.4 mL of 37°C base medium in a 1L sterile bottle. Pour off the old medium from the 10-stack cell culture flask and replace it with the transfection mixture (636mL). Incubate the cells for ~16 hours.

**Day 4:** Remove the transfection mixture from the 10-stack cell culture flask. Replace with 636 mL of 37°C base medium.

**Day 5:** Approximately 24 hours later collect the crude culture supernatant containing viral particles. Clarify by filtration using 250 mL 0.45μm stericups (Millipore). Up to four stericups might be necessary as the crude lysate clogs the filter past ~180 mL. At this step the crude lysate can be kept at 4°C for up to 24 hours. For concentrating the virus use a Beckman coulter ultracentrifuge with a swing bucket rotor (either SW28 or SW32Ti). In this protocol SW32Ti is used. Start by loading the rotor tubes under laminar flow. Insert thin wall polypropylene tubes into each rotor tube. Load 30 mL of crude lysate into each tube, securely close the caps. Use an analytic scale to confirm even distribution of weight for swinging buckets across from each other. Adjust volumes if necessary. Load the rotor with swinging buckets into the ultracentrifuge. Run a cycle of 19,500 rpm, 2 hours 32 minutes at 18°C. Repeat the run as many times as necessary (3x ≥). Discard the supernatant, re-suspend the pellet in 40 mL of 1 x PBS. Keep the concentrated sample at four degrees as the rest of it is processed. Incubate the concentrated virus on a rotor for 10 minutes. Freeze at 80°C until benzonase treatment.

**Benzonase treatment**

Benzonase activity 250 U/μL, working concentration 0.05 U/μL. Prepare a 7.5 U/μL benzonase stock solution in 2 mM MgCl in cell culture grade water. To make 1000 μL of 7.5 U/μL of benzonase, first prepare 1000 μL of 2 mM MgCl from the 1 M stock. Add 2 μL of 1 M MgCl (Millipor) into 998 $\mu$L of cell culture grade water. Remove 30 $\mu$L. Now make 7.5 U/$\mu$L benzonase stock in 2 mM MgCl. Add 30 $\mu$L of 250 U/$\mu$L benzonase into 2 mM MgCl. Final concentration is 0.05 U/$\mu$L, so the stock solution is now 150 x. Treat the concentrated viral lysates for 1 hour at room temperature, then aliquot 20$\mu$L per vial and freeze at -80°C.

**FASTA sequences of GFP and PIM-1 vectors**

>EG_001_CGW-PIM1-GFP_MND-seqPCR-FWD

NNNNNAGCGTGCGAACATCGATATCTTCTGGAGAGCAGTCACCATGCTCTCCCCA
GTGCATGCCCCAAGGACCTGAAATGACCCTGTGCCTTATTTGAACTAACCAATCAG
TTCGCTTCTCGCTTCTGTTCGCGCGCTTCTGCTCCCCGAGCTCTATATAAGCAGAG
CTCGTTTAGTGAACCGTCAGATCGCCTGGAGACGCCATCCACGCTGTTTTGACCTC
CATAGAAGATCAGTTAATTAAGAATTCCGGATGCTCTTGTCCAAAATCAACTCGCTT
GCCCACCTGCGCGCCGCGCCCTGCAACGACCTGCACGCCACCAAGCTGGCGCCC
GGCAAGGAGAAGGAGCCCCTGGAGTCGCAGTACCAGGTGGGCCCGCTACTGGGC
AGCGGCGGCTTCGGCTCGGTCTACTCAGGCATCCGCGTCTCCGACAACTTGCCG
GTGGCCATCAAACACGTGGAGAAGGACCGGATTTCCGACTGGGGAGAGCTGCCT
AATGGCACTCGAGTGCCCATGGAAGTGGTCCTGCTGAAGAAGGTGAGCTCGGGTT
TCTCCGGCGTCATTAGGCTCCTGGACTGGTTCGAGAGGCCCGACAGTTTCGTCCT
GATCCTGGAGAGGCCCGAGCCGGTGCAAGATCTCTTCGACTTCATCACGGAAAGG
GGAGCCCTGCAAGAGGAGCTGGCCCGCAGCTTCTTCTGGCAGGTGCTGGAGGCC
GTGCGGCACTGCCACAACTGCGGGGTGCTCCACCGCGACATCAAGGACGAAAAC
ATCCTTATCGACCTCAATCGCGGCGAGCTCAAGCTCATCGACTTCGGGTCGGGGG
CGCTGCTCAAGGACACCGTCTACACGGACTTCGATGGGACCCGAGTGTATAGCCC
TCCAGAGTGGATCCGCTACCATCGCTACCATGGCAGGTCGGCGGCAGTCTGGTCC
CTGGGGATCCTGCTGTATGATATGGTGTGTGGAGATATTCCTTTCGAGCATGACGA
AGAGATCATCAGGGGCCAGGTTTTCTTCAGCAGAGGGTCTCTTCAGGATGTCAGC
ATCTCATTAGATGGTGCTTGCCCTGAGACCATCAGATAGGCCAACCTTCGAAGAAA
TCCAGAACCATCCATGATGCAGATGTTCTCCTGCCCCAGGAACCTGCCTGAGAATC
CACCCTCCCCAGCCCTGNNGCCGGGGTCCCACCCANN

>EG_002_CGW-GFP_All-FP-FWD

CNGGGGGGAGGCGAGACTGTTCCCGGGGTGGTGCCCATCCTGGTCGAGCTGGAC
GGCGACGTAAACGGCCACAAGTTCAGCGTGTCCGGCGAGGGCGAGGGCGATGCC
ACCTACGGCAAGCTGACCCTGAAGTTCATCTGCACCACCGGCAAGCTGCCCGTGC
CCTGGCCCACCCTCGTGACCACCCTGACCTACGGCGTGCAGTGCTTCAGCCGCTA
CCCCGACCACATGAAGCAGCACGACTTCTTCAAGTCCGCCATGCCCGAAGGCTAC
GTCCAGGAGCGCACCATCTTCTTCAAGGACGACGGCAACTACAAGACCCGCGCCG
AGGTGAAGTTCGAGGGCGACACCCTGGTGAACCGCATCGAGCTGAAGGGCATCG

ACTTCAAGGAGGACGGCAACATCCTGGGGCACAAGCTGGAGTACAACTACAACAG
CCACAACGTCTATATCATGGCCGACAAGCAGAAGAACGGCATCAAGGTGAACTTC
AAGATCCGCCACAACATCGAGGACGGCAGCGTGCAGCTCGCCGACCACTACCAG
CAGAACACCCCCATCGGCGACGGCCCCGTGCTGCTGCCCGACAACCACTACCTG
AGCACCCAGTCCGCCCTGAGCAAAGACCCCAACGAGAAGCGCGATCACATGGTC
CTGCTGGAGTTCGTGACCGCCGCCGGGATCACTCTCGGCATGGACGAGCTGTACA
AGTAAAGCGGCCGCACTGTTCTCATCACATCATATCAAGGTTATATACCATCAATAT
TGCCACAGATGTTACTTAGCCTTTTAATATTTCTCTAATTTAGTGTATATGCAATGAT
AGTTCTCTGATTTCTGAGATTGAGTTTCTCATGTGTAATGATTATTTAGAGTTTCTCT
TTCATCTGTTCAAATTTTTGTCTAGTTTTATTTTTTACTGATTTGTAAGACTTCTTTTT
ATAATCTGCATATTACAATTCTCTTTACTGGGGTGTTGCAAATATTTTCTGTCATTCT
ATGGCCTGACTTTTCTTAATGGTTTTTTAATTTTAAAAATAAGTCTTAATATTCATGCA
ATCTAATTAACAATCTTTTTCTTTGTGGTT

### Supplemental figure legends

**Supplemental Figure 1. Soft agar assay.** (A) Positive control. HEK293T/17 (passage 5) cells 5,000/10,000/1,000 cells/well in triplicates after 10 days of incubation. (B) Colonies were counted using ImageJ software. $p^{***}<0.0001$, $p^{**}<0.001$ One-Way-ANOVA, Tukey's Multiple Comparison Test.

**Supplemental Figure 2. Cytochalasin B titration.** (A) control cCICs and (B) cCICs-PIM-1 from line H13-067 (p.11) treated with cytochalasin B. Incubation time 24 hours. Cytospin 400rpm for 3 minutes. Fixed in 4% PFA. Red=phalloidin. Blue=DAPI. 40x water objective.

**Supplemental Figure 3. Micronucleus Detection in HEK293T/17 Cells**. (A) HEK293T/17 cells as a positive control to demonstrate the sensitivity of the assay to capture micronuclei.

**Supplemental Figure 4. PIM-1 transfection selectively increases transcription of PIM-1 variant 1.** PIM-1 transfection selectively increases transcription of PIM-1 variant 1, as revealed by (A) detected PIM-1 Fragments Per Kilobase Million (FPKM) per group and (B) spliced alignment to genomic position.

**Supplemental Figure 5. PIM-1 transfection selectively increases transcription of PIM-1 variant 1 in patient derived cell lines.** A) PIM-1 is upregulated in splice variant 1 (bottom row) but not in variant 2 (middle row) or 3 (top row) in PIM-1 overexpressed cCICs.

**Supplemental Figure 6**. **PIM-1 overexpression minimally impacts the transcriptome.** (A) PIM-1 enhancement minimally impacts the overall transcriptome and (B) its relationship with differentially expressed genes (FPKM>0). (C) PIM-1 enhancement minimally impacts the high-abundance expressed genes and (D) its relationship with differentially expressed genes (FPKM >1).

**Supplemental Figure 7**. **PIM-1 and PIM-2 kinase activity revealed by real-time fluorescent biosensor quantitation.** (A) Progress curves with the AQT57218 1C09 sensor for PIM-1, 2 and 3 and several off-target kinases (CAMK and AGC group) known to phosphorylate PIM-1 substrates and (B) Histogram of initial reaction rates for these 8 kinases. Data demonstrate that the AQT57218 1C09 sensor preferentially detects the activity of PIM-1 and PIM-3, with much less activity of PIM-2 and no reactivity with 6 other kinases from either the CAMK (CHEK2, MK2 also known as MAPKAPK2) or AGC (AKT3, p70S6K and RSK1) groups, despite the fact that the PIM kinases belong to the same CAMK group and share certain substrates with AGC group kinases.

**Supplemental Figure 1. Soft Agar Assay**

A.

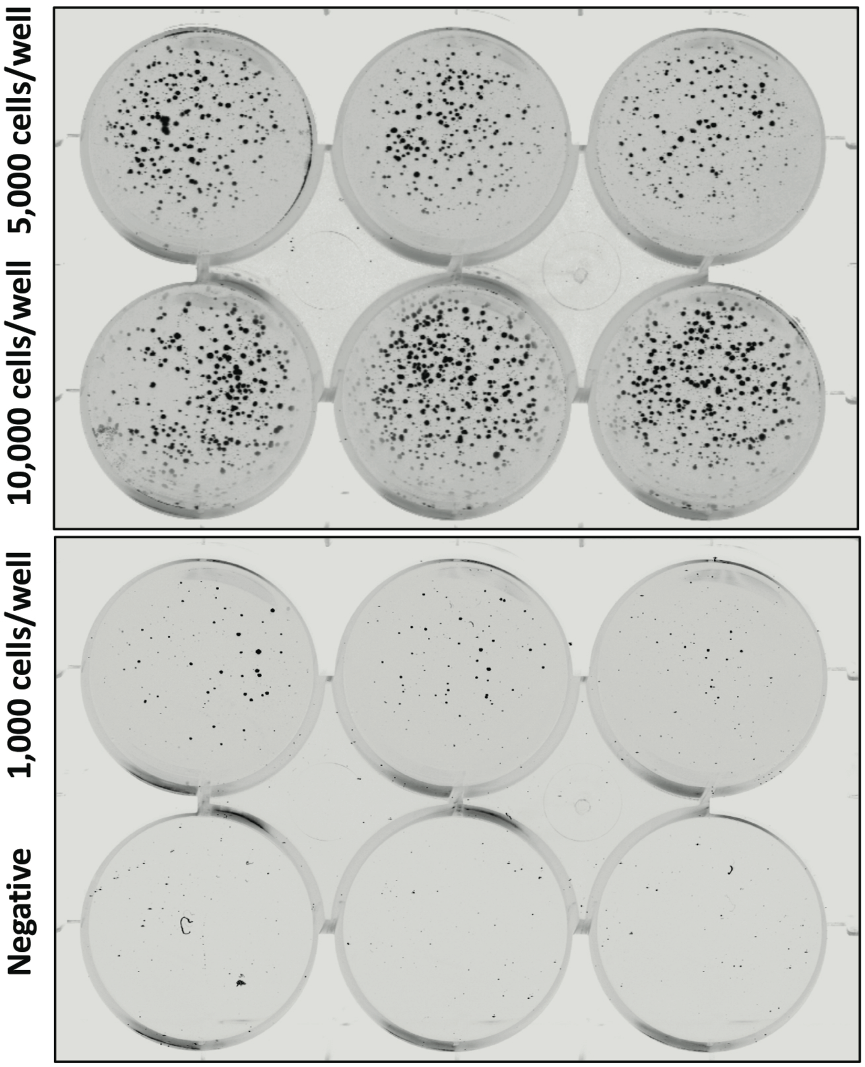

B.

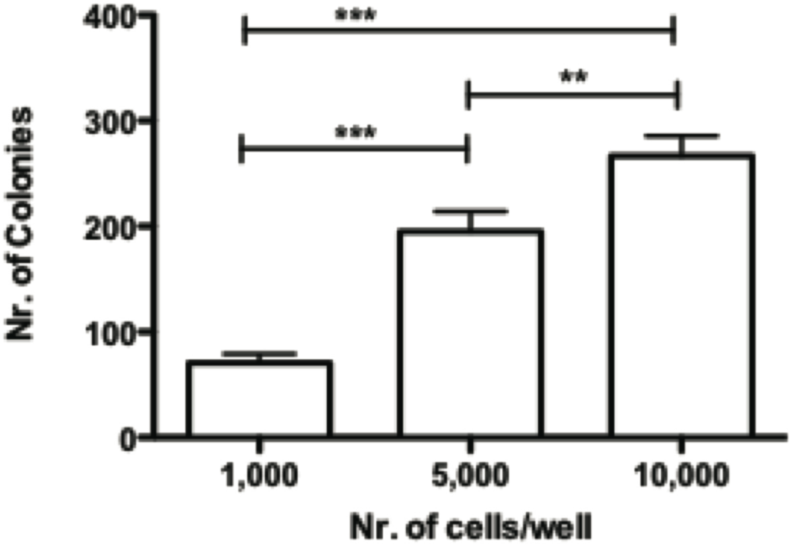

**Supplemental Figure 2. Cytochalasin B titration.**

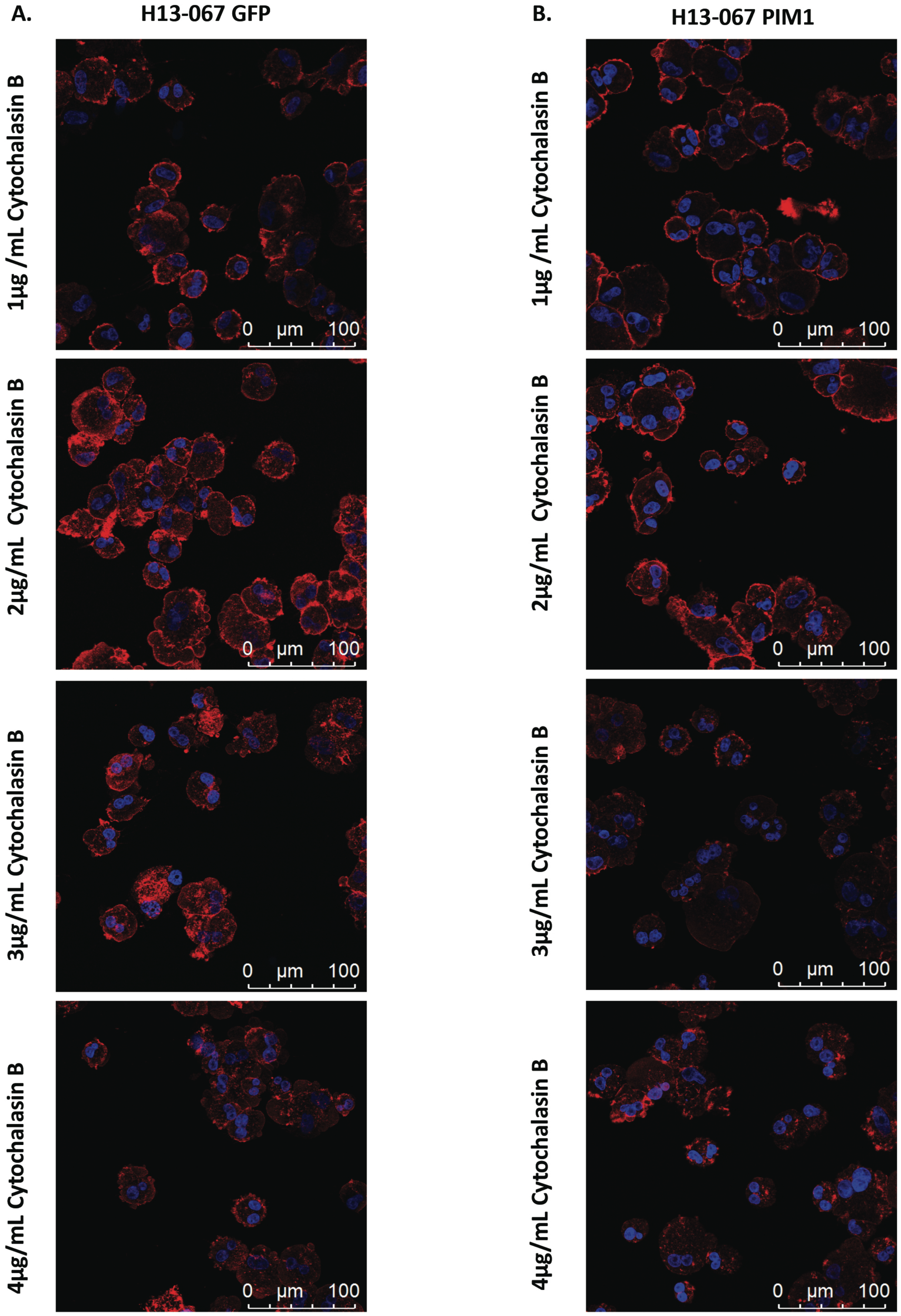

**Supplemental Figure 3. Micronucleus Detection in HEK293T/17 Cells.**

**A.**

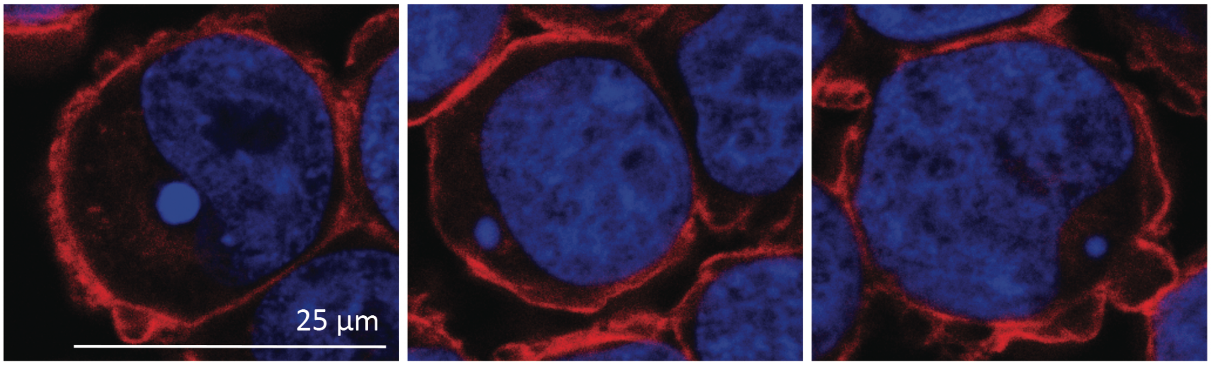

**Supplemental Figure 4. PIM-1 transfection selectively increases transcription of PIM-1 variant 1.**

**A.**

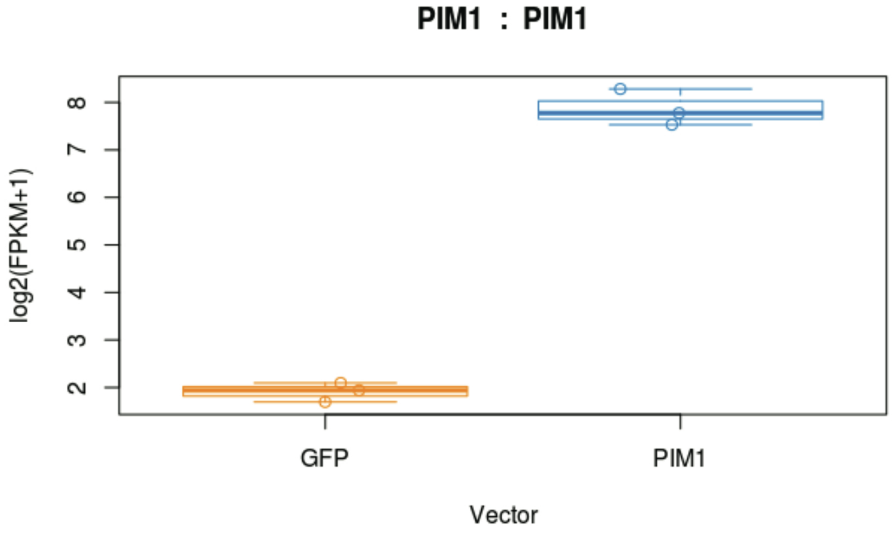

**B.**

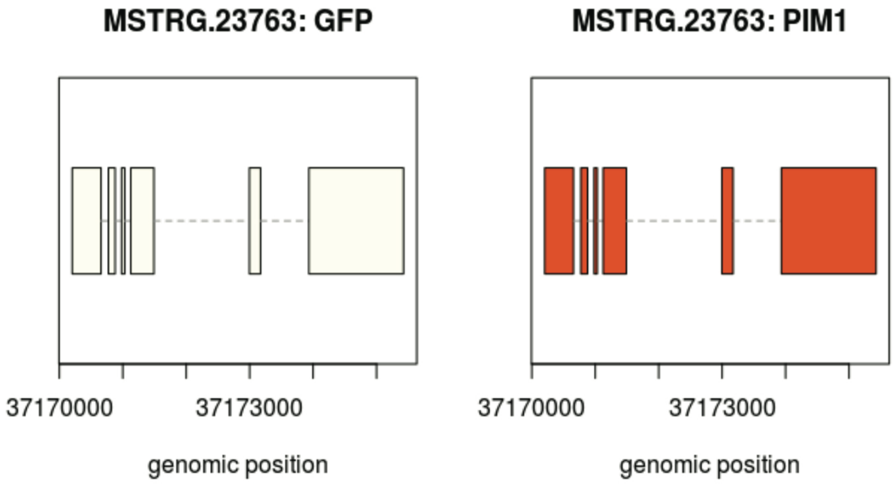

**Supplemental Figure 5. PIM-1 transfection selectively increases transcription of PIM-1 variant 1 in patient derived cell lines.**

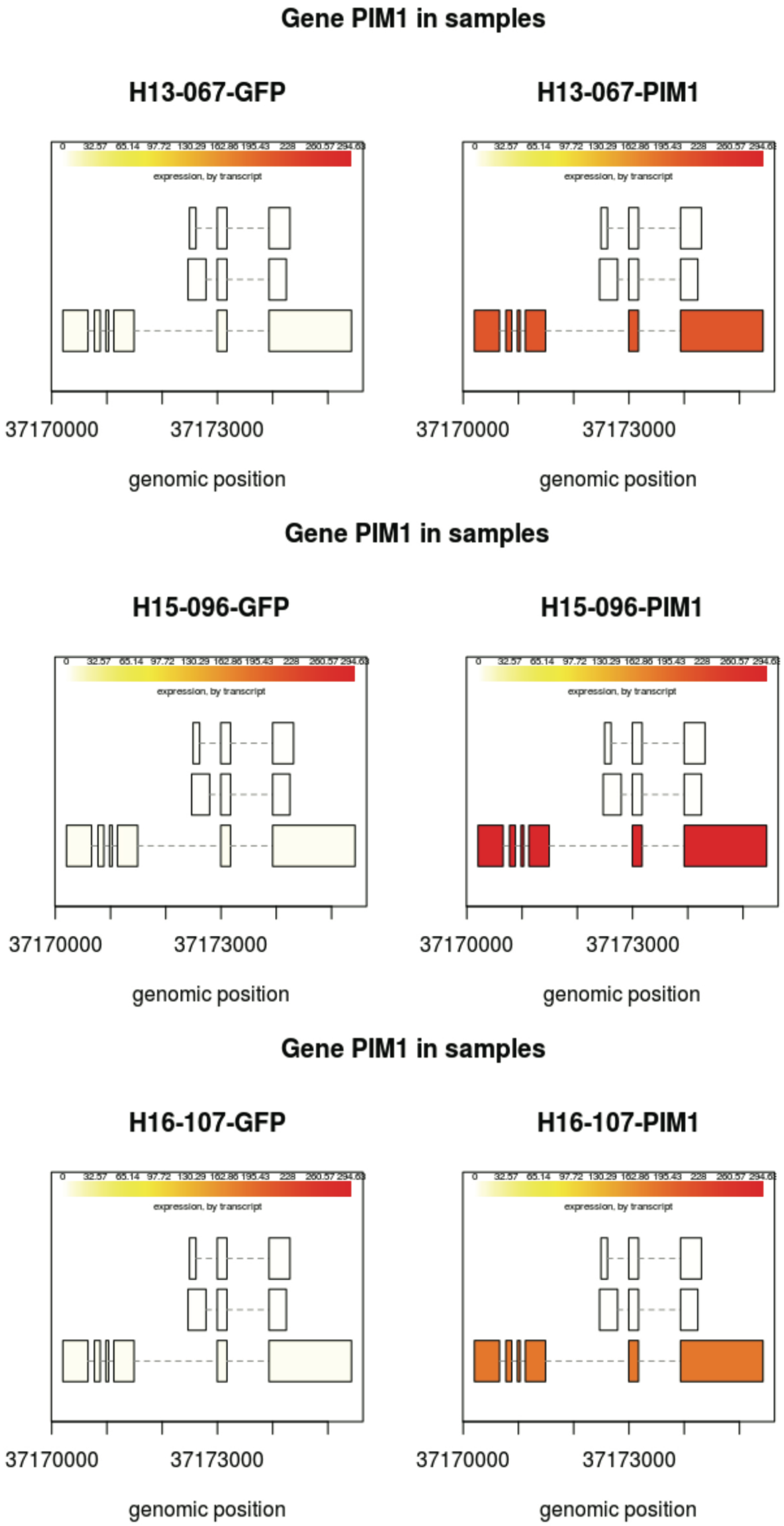

**Supplemental Figure 6. PIM-1 overexpression minimally impacts the transcriptome.**

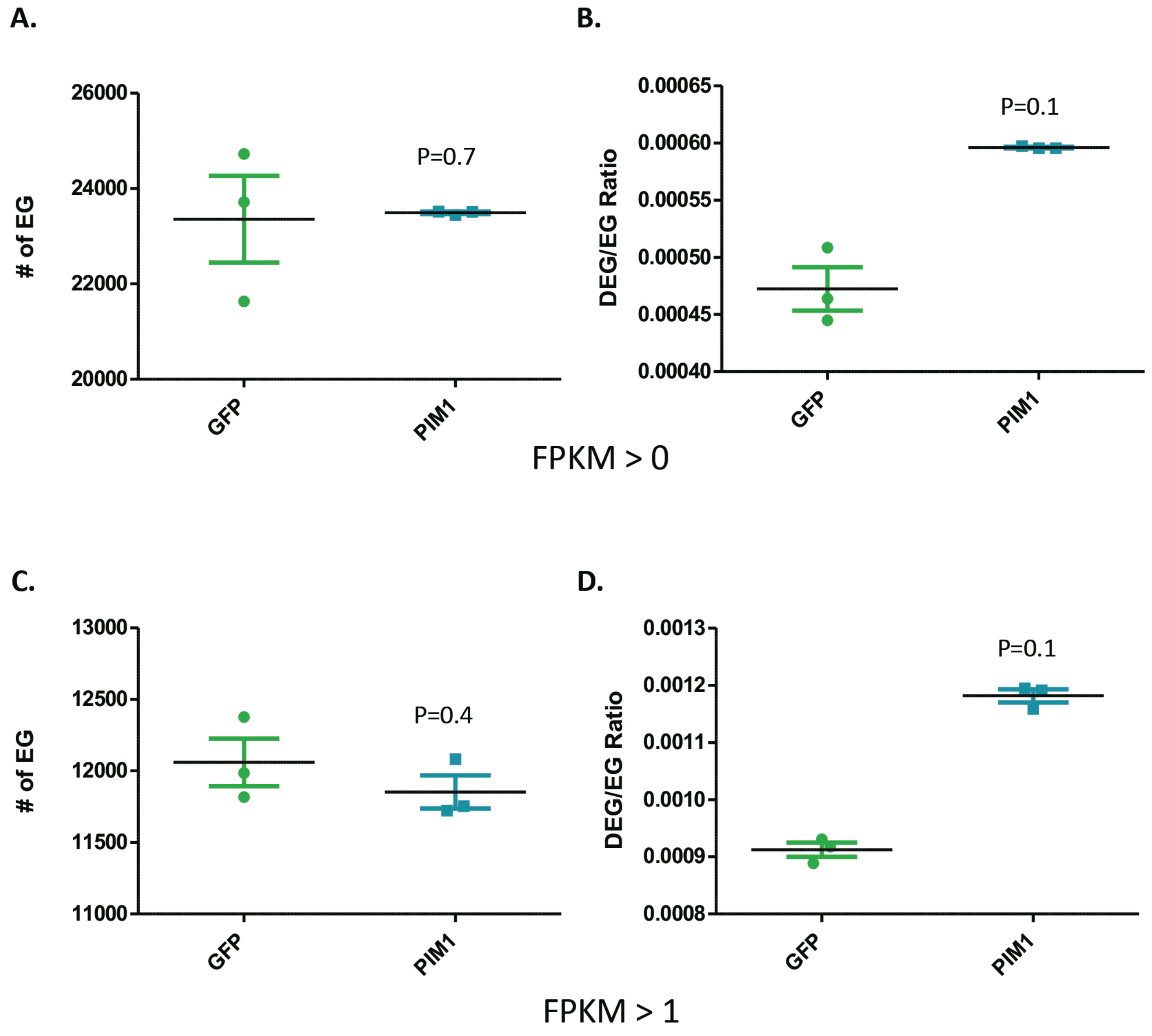

**Supplemental Figure 7. PIM-1 and PIM-2 kinase activity revealed by real-time fluorescent biosensor quantitation.**

**A.**

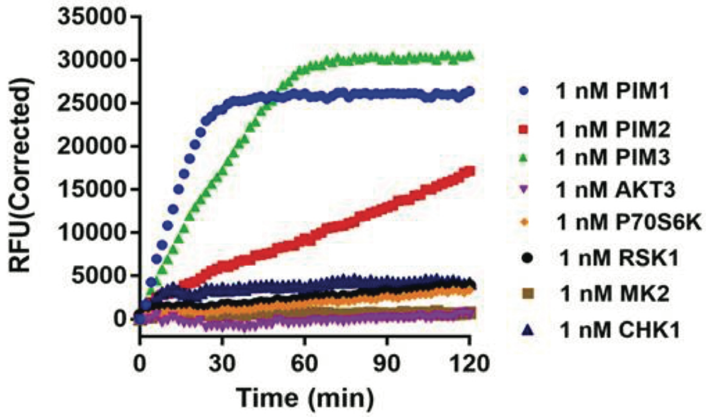

**B.**

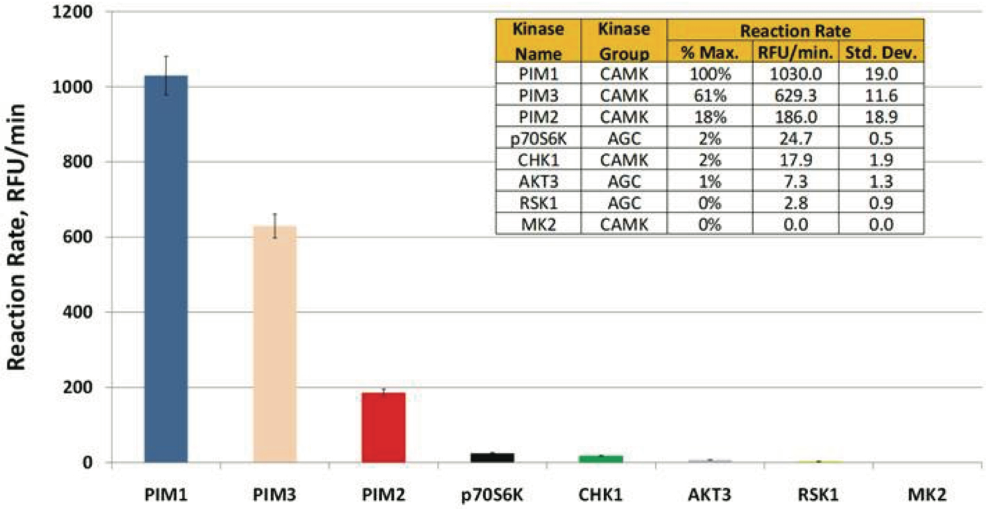

| Kinase Name | Kinase Group | Reaction Rate % Max. | RFU/min. | Std. Dev. |
|---|---|---|---|---|
| PIM1 | CAMK | 100% | 1030.0 | 19.0 |
| PIM3 | CAMK | 61% | 629.3 | 11.6 |
| PIM2 | CAMK | 18% | 186.0 | 18.9 |
| p70S6K | AGC | 2% | 24.7 | 0.5 |
| CHK1 | CAMK | 2% | 17.9 | 1.9 |
| AKT3 | AGC | 1% | 7.3 | 1.3 |
| RSK1 | AGC | 0% | 2.8 | 0.9 |
| MK2 | CAMK | 0% | 0.0 | 0.0 |

**Supplemental Table 1.**

| | Component | Catalog Number |
|---|---|---|
| Human ckit+ Cardiac Interstitial Cell Medium | F12 HAM's (1x) | SH30026.01, HyClone |
| | 10% ES FBS | 16141061, Gibco |
| | 1% Penicillin-Streptomycin-Glutamine (100X) | 10378016, Gibco |
| | 5mU/mL Human Erythropoietin | H5166, Sigma |
| | 10ng/mL Recombinant Human FGF-basic | GMP100-18B, Peprotech |
| | 0.2mM L-Glutathione | 10007461, Cayman |
| HEK293T/17 Medium | DMEM (1x) | 11995-040, Gibco |
| | 10% FBS | FB-01, Omega Scientific |
| | 1% Penicillin-Streptomycin-Glutamine (100X) | 10378016, Gibco |
| (2x) Human ckit+ Cardiac Interstitial Cell Medium | Cell Culture Grade Water | SH30529.LS, HyClone |
| | F-12 Nutrient Mixture (Ham) | 21700-075, Gibco |
| | 20% ES FBS | 16141061, Gibco |
| | 2% Penicillin-Streptomycin-Glutamine (100X) | 10378016, Gibco |
| | 10mU/mL Human Erythropoietin | H5166, Sigma |
| | 20ng/mL Recombinant Human FGF-basic | GMP100-18B, Peprotech |
| | 0.4mM L-Glutathione | 10007461, Cayman |
| | 2.4g/L $NaHCO_3$ | S5761, Sigma |
| (2x) HEK293T/17 Medium | Cell Culture Grade Water | SH30529.LS, HyClone |
| | DMEM powder, high glucose | 12100061, Gibco |
| | 20% FBS | FB-01, Omega Scientific |
| | 2% Penicillin-Streptomycin-Glutamine (100X) | 10378016, Gibco |
| | 7.4 g/L $NaHCO_3$ | S5761, Sigma |

**Supplemental Table 2.**

| | | | | | | | | |
|---|---|---|---|---|---|---|---|---|
| Including mock plate (+75 $cm^2$) | | | | | | | | |
| DMEM with no adds for 10-Stack (6360 $cm^2$ + 75 $cm^2$ = 6435) | | | | | **21.2** | **mL** | | |
| PEI for 10-Stack (6360 + 75 $cm^2$) | | | | | **5.3** | **mL** | | |
| For 10-Stack (6360 + 75$cm^2$) | | | | | | | | |
| | | | | DMEM | **105.8** | **mL** | | |
| **Plasmids** | **µg for 6435 $cm^2$** | | **Stock [c]** | | **Volume (uL)** | | **Volume (mL)** | |
| pVSVG | 381.0 | µg | 1.4 | µg/µL | **272.2** | **µL** | **0.3** | **mL** |
| pRev | 264.6 | µg | 1 | µg/µL | **264.6** | **µL** | **0.3** | **mL** |
| pGal/Pol | 529.2 | µg | 1.4 | µg/µL | **378.0** | **µL** | **0.4** | **mL** |
| GOI | | | | | | | | |
| CGW PIM1 GFP | 1058.4 | µg | 0.6 | µg/µL | **1764.0** | **µL** | **1.8** | **mL** |
| CGW GFP | 1058.4 | µg | 0.9 | µg/µL | **1176.0** | **µL** | **1.2** | **mL** |
| **Add all plasmids together in a 150 ml sterile bottle, and then divide into 3 equal parts.** | | | | | | | | |
| **Plasmid mixture total volume** | | **109.7** | **mL** | | | | | |
| **Into each 50 mL tube** | | **36.6** | **mL** | | | | | |
| **PEI mixture into each 50 mL tube** | | **8.8** | **mL** | | | | | |
| **Total volume** | **136.2** | **mL** | | | | | | |
| **Plasmid mix for 75 $cm^2$** | **1.6** | **mL** | | | | | | |
| **Medium for 6360 $cm^2$** | **501.4** | **mL** | | | | | | |
| **Medium for 75 $cm^2$** | **5.9** | **mL** | | | | | | |
